# Multi-model reinforcement learning with online retrospective change point detection

**DOI:** 10.1101/2025.05.13.653727

**Authors:** Augustin Chartouny, Mehdi Khamassi, Benoît Girard

## Abstract

Humans continuously adapt to uncertain and changing situations. However, most reinforcement learning models of human behavior struggle to explain this capability. We propose a novel reinforcement learning agent for uncertain and volatile Markov decision processes, which we call Multi-Model with Retrospective Change Point Detection (MMRCPD). MMRCPD relies on two novel ideas: arbitrating between local models rather than contexts of the environment and retrospectively detecting change points. Arbitrating between local models limits memory costs and enables faster adaptation to new contexts which sub-parts have been experienced before. Retrospective change point detection mimics the capacity of humans to infer the latent cause of a change after it happened and maintain precise models of the environment. MMRCPD can detect local changes online, create new models, retrospectively update its models based on when it estimates that the change happened, reuse past models, merge models if they become similar, and forget unused models. This novel multi-model agent outperforms single-model and context-level change-detection methods in uncertain and locally changing environments. These results yield new insights and predictions concerning optimal decision-making in changing and uncertain environments, which could in turn be tested in behavioral experiments.

## 1 Introduction

Humans constantly adapt to uncertain, new, and open-ended environments in their daily lives. To explain this behavioral flexibility, researchers suppose that humans perform reinforcement learning (RL) and contextual inference. Standard RL explores how humans, animals, or artificial agents learn and refine their strategies over time to make decisions based on their interactions with an environment [Sutton and Barto, 2018]. Contextual inference, on the other hand, refers to the online interpretation of environmental cues to discriminate between different situations [Monsell, 2003]. Modeling how humans infer contexts and refine their strategies based on the inferred context is essential to describe human behavior.

Traditional approaches to contextual inference in RL [Hadoux et al., 2014], decision-making [Collins and Koechlin, 2012] and robotics [Caluwaerts et al., 2012] study changes at the whole-task level. When detecting a change, these context-level approaches change all of their beliefs about the environment simultaneously. However, the assumption that humans, animals or artificial agents differentiate task changes solely at the context level does not facilitate the transfer of knowledge between tasks that share common properties. Context-based models relearn an entire context to adapt for minor variations, which is sometimes inefficient [Khetarpal et al., 2022]. For instance, a rat exploring a maze is not obligated to relearn the entire structure of the maze every time a door is opened or closed [Martinet et al., 2011]. As the context only changes locally, the rat is only required to learn two models for the door, and can use a shared model for the rest of the maze.

In addition, detecting a task change at the context-level often relies on aggregate measures such as cumulative sums of local changes [Hadoux et al., 2014]. Thus, contextual inference models often have trouble identifying the exact factors that drive contextual shifts, including the precise moment or location of each change. This, sometimes added with high computational costs [Gershman et al., 2010], makes it impossible to retrospectively assign observations to the correct context after a change has been identified. As a result, context-level approaches tend to assume that a change occurs just before or at the exact point of detection [Collins and Koechlin, 2012, Da Silva et al., 2006, Hadoux et al., 2014]. In contrast, humans make sense of a detected change by retrospectively inferring a latent cause [Rensink, 2002, Moran et al., 2019]. Context-level methods that lack the ability to retrospectively assign observations to the correct context fail to capture this fundamental aspect of change detection in humans.

To address local changes and retrospectively detect when they happened, we propose a novel model-based RL agent, which we call Multi-Model with Retrospective Change Point Detection (MMRCPD). MMRCPD creates local models for each local change and switches between them based on environmental observations. This method enhances the transfer of knowledge between contexts and maintains accurate models of the environment, using retrospective change point detection.

### 1.1 Context- and local-level approaches

The definition of contexts varies across decision-making and RL domains. Contexts are generally considered to be an ensemble of properties of the environment that an agent must perceive to behave efficiently [Heald et al., 2023]. We consider here that a context is a mode of a Markov Decision Process, using the formalism of Hidden-Mode Markov Decision Processes [Choi et al., 1999]. The environment the agent interacts with can be in different stationary Markov Decision Processes, namely modes or contexts, and the agent has no control over when the environment changes from one context to another. Each context is likely to remain stable for some time, and we make no assumptions regarding the total number of contexts an agent may face. This is similar to the unbounded number of latent causes from the Bayesian literature [Gershman et al., 2010]. An example environment with different contexts (Figure 1, Top) is a non-stationary three-armed bandit task [Wu et al., 2018, Cinotti et al., 2019]. For a three-armed bandit, a context is comprised of three reward distributions – one for each of the three arms. The context changes when the distribution of one or several arms change [Beaumont et al., 2025].

**Figure 1:**
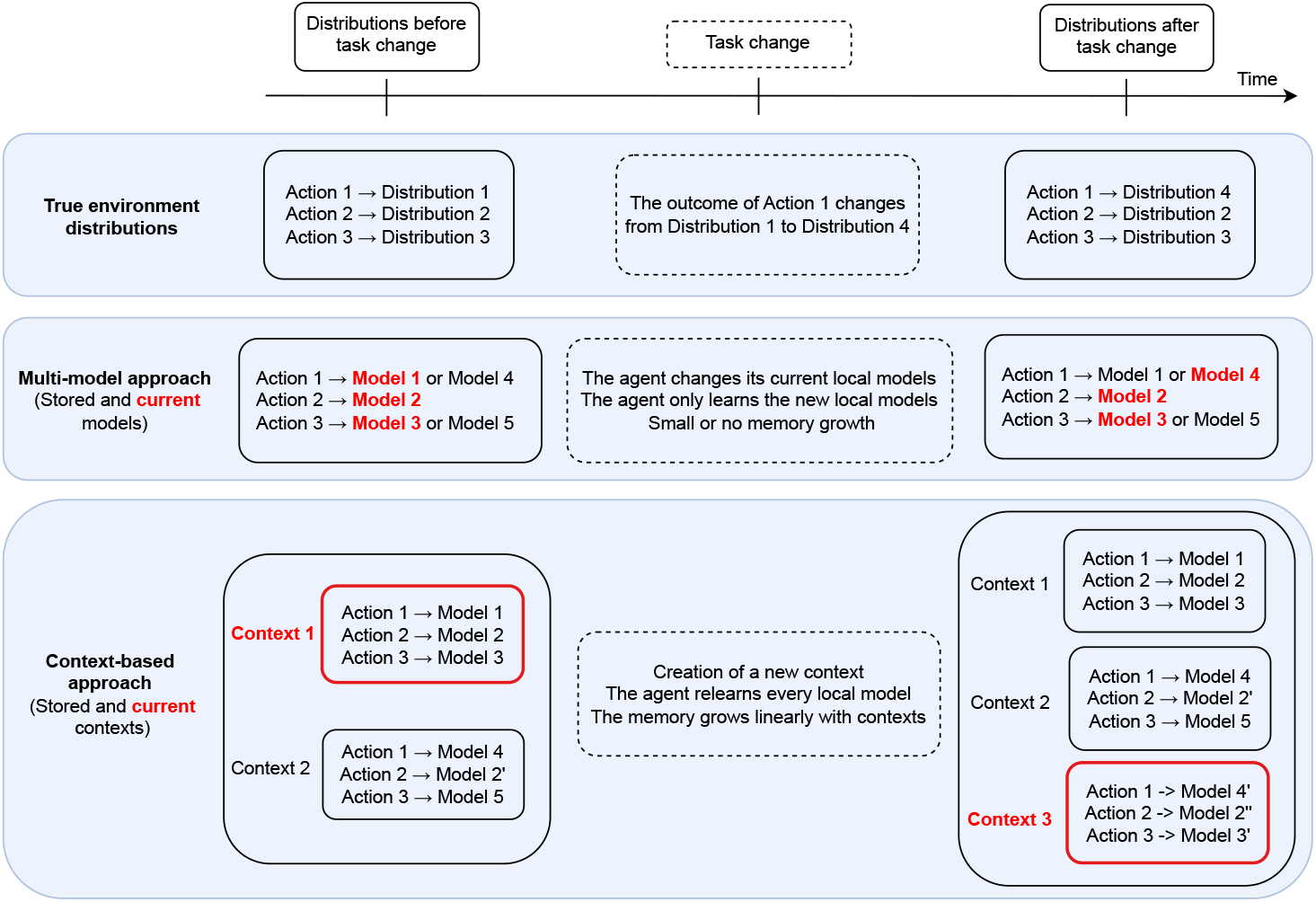
Comparison between context-level and local-multi-model approaches in a three-action task. Both approaches model the result of each action, for example the distribution of rewards in a three-armed bandit. If contexts share features, arbitrating directly between models is theoretically more efficient than arbitrating between contexts. The task change involves a change in the result of action 1 from distribution 1 to distribution 4. The context-level approach must create a new context and relearn a model for each action. The models the context-level agent relearn are indicated with apostrophes. For example, models 2, 2′, and 2′′ are three different models of distribution 2. In contrast, the multi-model approach only relearns the local models that changed during the task change. Red text indicates the best context or the best models for each method based on the true environment distributions.

Context-level methods modify all local models learned when detecting a context change [Da Silva et al., 2006, Hadoux et al., 2014, Collins and Koechlin, 2012]. As mentioned above, this grants prompt adaptation to changes only after an agent has learned all possible contexts, thus preventing the leveraging of cross-context similarities when a new context is being learned. Moreover, memory usage increases linearly as a function of the number of contexts, irrespective of cross-context similarities (Figure 1, Bottom). Context-level approaches may also be inefficient in detecting task changes when an environment is only modified locally, due to the high degree of similarity between contexts. Subsequently, if local changes occur frequently, context-level approaches may fail to adapt or require full context updates consistently. Both of these issues would increase computational costs and reduce performance.

Considering the above-outlined disadvantages of context-level approaches, we believe that a local-model-level approach would be more appropriate to detect changes between contexts that share similar features. We propose an alternative approach that detects changes in local models (as opposed to entire contexts), adapting to them accordingly (Figure 1, Center). Beyond reducing memory usage and computational costs, this approach favors transfer learning; Only cross-context differences are learned and stored. The local-model-level approach encourages adaptation to local changes and promotes the transfer of prior knowledge to new contexts.

### 1.2 Retrospective change point detection

Detecting changes at the local level facilitates finding the source of surprising events, which may come from the uncertainty of a stable environment (stochasticity) or a change in the environment (volatility). To categorize surprising events from an observer’s perspective, the decision-making literature often disentangles expected uncertainty from unexpected uncertainty [Yu and Dayan, 2005]. Expected uncertainty consists of the predicted variability of outcomes in a stable environment. Unexpected uncertainty consists of events that violate model expectations and can come from a change in the environment. To adapt to uncertain (stochastic) and changing (volatile) environments, model-based decision-makers must assess to what extent each observation modifies their beliefs about the environment. Surprising observations decrease the reliability of the learned model and sometimes induce stronger updates [Nassar et al., 2010].

However, one surprising event may not be enough to assess the reliability of a model. For example, when trying to understand whether a die is fair or rigged, observing one roll with an outcome between 1 and 6 is not very informative. Conversely, observing only sixes for the first five rolls indicates that the die is rigged. To further understand how to detect a change in a distribution, we consider the toy problem of a passive observer trying to determine whether a die is rigged or fair. The observer only has access to the results of the rolls. Nevertheless, they know that there are only two dice, one fair and one rigged. Moreover, they understand that the chance to switch from one die to another is small after a die change and that each die is rolled several times in a row. The roll sequence is the following, with labels indicating rigged or fair die for the reader:

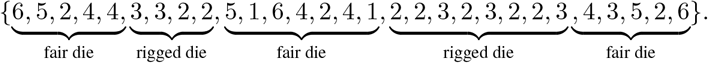

Depending on whether the agent receives the roll results sequentially or can access the entire sequence directly, their prediction of what die was rolled at each time step should change. For example, since the probability distributions of the two dice overlap, the agent can only detect surprising sequences a few trials after a die change. This means that once the agent detects a change, it should retrospectively estimate the moment the task changed, before updating its models for each die.

Moreover, even if the agent observes two 2s and two 3s the first time it faces the rigged die, they cannot know whether this was an unlikely sequence with the fair die or if the die changed. However, when the agent observes the second sequence of 2s and 3s, they can create a new model for the second die, as this sequence is highly unlikely with a fair die. With this new model of the rigged die, the agent will detect future die changes more quickly. When looking back at the whole sequence with an accurate model of the rigged die, the agent may predict that the first sequence of 2s and 3s came from the rigged die, even though it did not detect the change at the time.

From this illustrative example, we extract three pivotal ideas for the model discrimination we will use. (1) An agent needs several observations to detect a task change and should not create a new model based on a single surprising observation. (2) An agent needs to be able to retrospectively detect a change point, to reassign the observations to the correct models. (3) An agent with an infinite memory could detect prior changes it missed by learning the approximate dynamics of the various models. However, storing long sequences of observations would induce memory costs. One possible solution is to work with finite-observation windows and create more models than needed in ambiguous cases, even if it means merging some models later on.

### 1.3 Contributions

We present a new multi-model RL agent which adapts to uncertain and changing environments. This agent disentangles stochasticity from volatility by maintaining several models of the environment. Contrary to previous work, our approach arbitrates between local models instead of contexts. This local-model-level method allows for the transfer of knowledge between different contexts with shared sub-parts, while being more memory efficient than context-level methods. We also introduce a new technique using negative log-likelihood minimization to accurately detect when changes occur. We demonstrate that our multi-model agent outperforms single-context and multi-contexts RL agents in locally changing uncertain environments. Additionally, the multi-model approach we introduce has limited computational cost and is stable to parameter changes.

## 2 Multi-Model with Retrospective Change Point Detection

### 2.1 General description of MMRCPD

MMRCPD is a change detection agent which follows a decision-tree at each time step to adapt to uncertain and changing environments (Figure 2). We can decompose this agent in several modules with distinct functions. **Learning**: After performing an action in the environment, MMRCPD updates its model based on the observed outcome and classical reinforcement learning formulas. **Change detection**: At each time step, MMRCPD checks if its current local model adequately explains the latest local observations. If a stored model explains the last observations better, or if no model explains the last observations well, the agent changes its current local model. **Change point finding**: When MMRCPD detects a change, it retrospectively computes the change point, finds the associated new local model, and reassigns the observations after this change point to the new model. This new model becomes the current model and the agent starts using it to solve the task. **Merging:** At every step, if two models are similar, the agent merges them. This limits computational costs by reducing the number of unnecessary computations, and to be less conservative with model creation. **Forgetting:** If the memory storage of the agent exceeds a preset threshold, it forgets an unused model.

**Figure 2:**
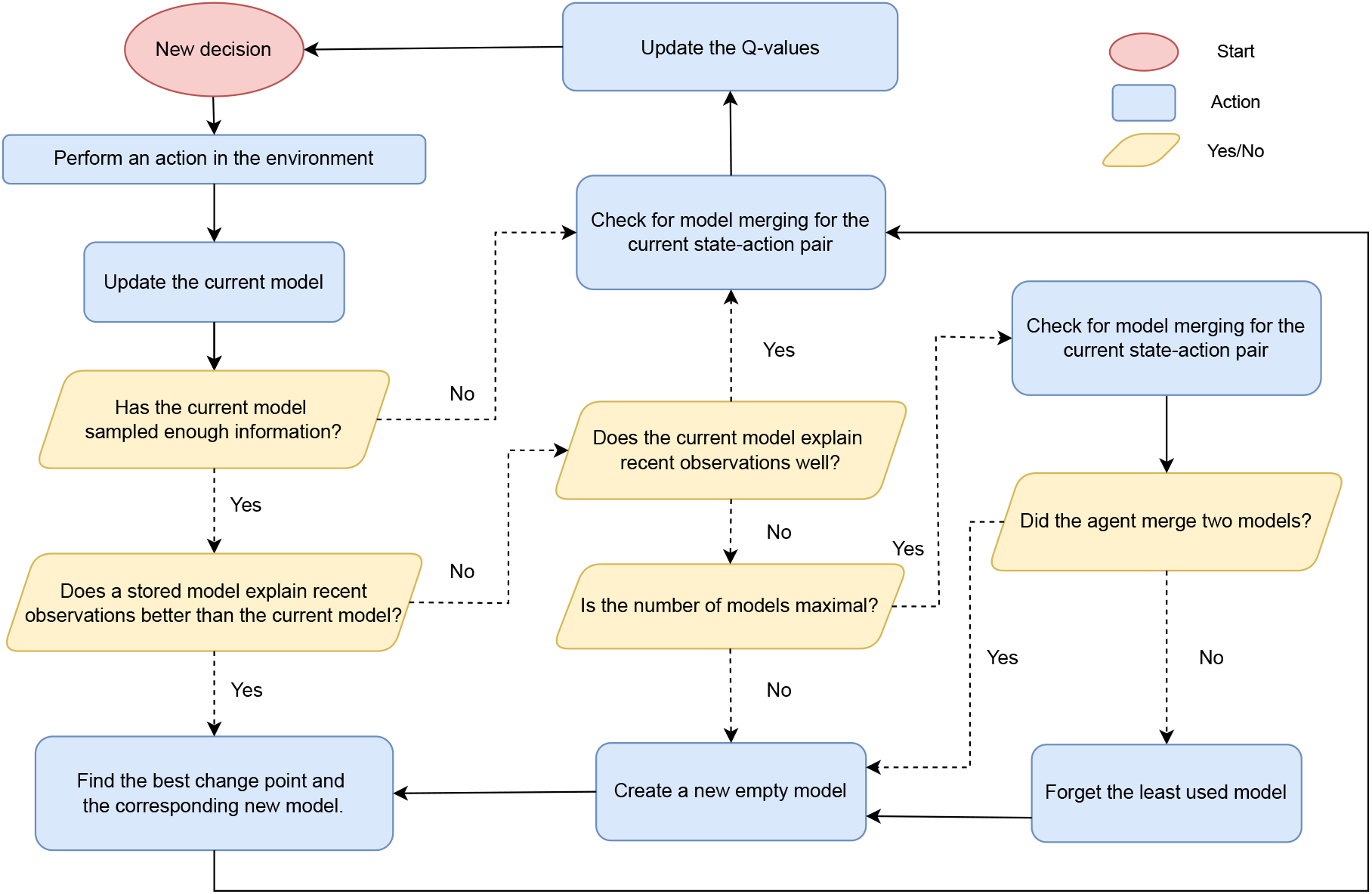
Structure of MMRCPD, presented as a decision tree. Based on yes/no tests, the tree describes what the agent does for each new decision (top-left red circle). For example, if a stored model explains recent observations better than the current model, the agent finds the best change point and changes model with the corresponding new model.

### 2.2 Model learning

#### 2.2.1 Learning models in RL

We consider the general case of a RL agent acting in a Markov decision process {*S, A, T, R*}. The agent has access to a set of states *S* and of actions *A. T* is a transition function that maps state-action pairs to the conditional probability distribution on the possible arrival states and *R* is a reward function mapping the state-action pairs to a distribution of real values.

##### Single-model infinite-horizon approach

Classical model-based RL agents learn an internal model of the environment for each state-action pair by sampling tuples of observations [Sutton and Barto, 2018]. For each state-action pair (*s, a*), we define the reward model *M*_*R*_(*s, a*) and the transition model *M*_*T*_ (*s, a*) as the set of rewards and transitions the agent has observed from performing action *a* in state *s*. Given (*s*_*t*_, *a*_*t*_, *r*_*t*_) as the state, the action, and the immediate reward at time *t*, we denote

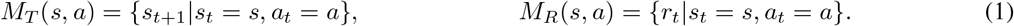

An infinite-horizon model-based agent tries to estimate the true transition function *T* and reward function *R* based on all the observations it has sampled. A simple approach to approximate *T* and *R* is to compute the frequency of arrival states 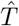 and the mean reward 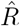. The agent then uses 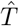 and 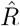 to plan and decide which action to take, using methods from dynamic programming such as value iteration [Sutton and Barto, 2018]. With 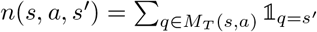 the number of times the agent arrived in state *s*′ after making action *a* in state *s* and *n*(*s, a*) = | *M*_*T*_ (*s, a*) | = | *M*_*R*_(*s, a*)) | the number of times the agent took action *a* in state *s*,

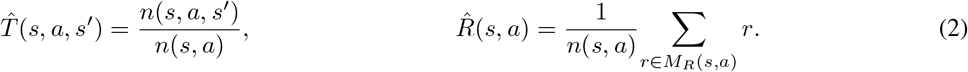

##### Single-model finite-horizon approach

A common approach to deal with non-stationary environments is to consider only the last *h* observations for the transition and the reward models [Massi et al., 2022]. The agent forgets the oldest observation of a model when the number of observations in the model exceeds *h*. We define 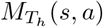 and 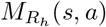 as the *h* last rewards and arrival states the agent sampled for (*s, a*). If the agent sampled less than *h* observations, then 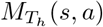 and 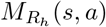 correspond to *M*_*T*_ (*s, a*) and *M*_*R*_(*s, a*) from Equation 1. If the agent sampled *h* observations or more, we can write 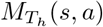 and 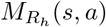 as

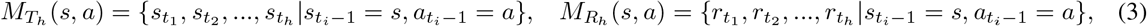

where *t*_1_ *< t*_2_ *<* … *< t*_*h*_ are the most recent *h* time steps for which 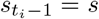 and 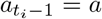. The associated count functions are 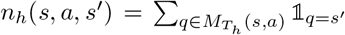 and 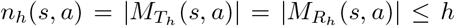. The frequentist finite-horizon transition functions 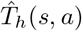 and reward function 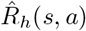 become

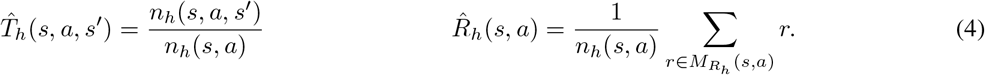

Finite-horizon approaches update their models faster than infinite-horizon approaches, as they replace old observations with new ones. However, the precision of their models is limited by the fixed number of recent observations *h* they consider. For example, if the true reward and transition functions are uncertain, *h* = 10 observations may not be enough to describe them. A finite-horizon model may also be unstable with low values of *h*, as each new observation accounts for 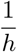 of the estimated distribution. Finally, finite-horizon agents optimize new tasks as they come but they cannot re-use previous observations or learn multiple tasks simultaneously.

##### Multi-model approach

Another possible approach to adapt to task changes is to arbitrate between multiple competing models with infinite horizons. In this multi-model approach, the agent stores several transition and reward models and changes its current models depending on the situation it faces. The agent associates each observed arrival state and reward to one of its transition or reward models. We define 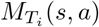 and 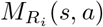 as the ensemble of observations that have been attributed to the transition model *i* and the reward model *i*. We define *i*_*T*_ ′(*t*) and *i*_*R*_′(*t*) as the models the agent uses for (*s, a*) at time *t*. When making action *a* in state *s*, the multi-model attributes the new observations to its current reward and transition models. Hence, 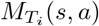 and 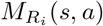 are

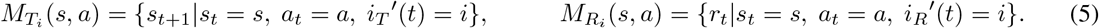

Each model has an infinite memory and the agent can use Equation 2 to compute an approximate transition function 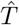 and reward function 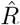.

#### 2.2.2 Implementation of model learning

MMRCPD uses a multi-model approach. At each time step, the agent adds the observed reward and arrival state to its current reward model and transition model, while the other reward and transition models do not change (Equation 5). MMRCPD records the last *h* arrival states and rewards received observed by making action *a* in state *s*. It also keeps track of the local model used at the time of each observation. Depending on how closely the agent’s transition model or reward model aligns with the observed data of the last *h* steps for (*s, a*), the agent either continues updating the current local model, changes to a stored model or creates a new one. We present the methods and results for MMRCPD when it arbitrates between local transition models. However, these same methods can also be applied to local reward models, provided the agent learns a distributional reward function.

### 2.3 Detecting a change

#### 2.3.1 How to evaluate model reliability with incoming observations?

The goal of the change detection module is to find when the current model no longer explains recent observations. Classical approaches to measure if a distribution explains incoming data is to use statistical tests or information theory methods. In decision-making, many change detection methods use Bayesian inference to compute an estimated hazard rate or detect a change point online [Yu and Dayan, 2005, Gershman et al., 2010, Wilson and Niv, 2012, Collins and Koechlin, 2012]. When not maintaining Bayesian posteriors, methods from the reinforcement learning literature often use sequential and incremental change detection methods, such as cumulative sums [Hadoux et al., 2014]. To assess the reliability of its models, MMRCPD directly compares them with the distribution of the recent observations using a Kullback-Leibler (KL) divergence. This formulation takes inspiration from Bayesian surprise [Itti and Baldi, 2009], which assesses to what extent one observation modifies the belief about a model. The choice of the KL divergence is motivated by the asymmetry between the recent observations, which are limited and variable, and the learned models which are generally larger and more stable. MMRCPD uses this asymmetry, by putting the distribution of recent observations first and stored models second in the computation of the KL divergence.

#### 2.3.2 Implementation of change detection

At each time step, MMRCPD verifies if the current model explains most recent observations well. If a stored model explains the recent observations better than the current model, or if the current model fails to explain the current observations, the agent changes its current model.

We suppose that for a given state and action (*s, a*), the agent stored *k* models 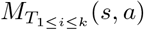 and that the current model is model *i*′. Every time the agent takes action *a* in *s*, it computes the Kullback-Leibler divergence between each stored model and the distribution of the last *h*_*i*_*′* observations, with *h*_*i*_*′* being the last consecutive observations attributed to the current model *i*′ in the last *h* passages. Thus *h*_*i*_*′* ≤ *h* and *h*_*i*_*′* = *h* if the last *h* passages were attributed to model *i*′. The agent computes the Kullback-Leibler divergence between the distribution of the last observations 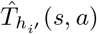 and the distribution of each stored model 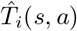 (Equation 6).

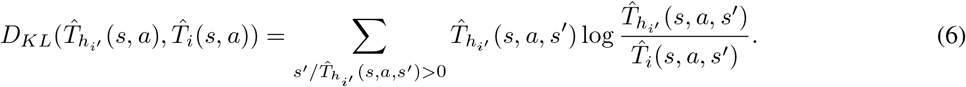

If the support of the first distribution 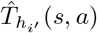 is not included in *T*_*i*_(*s, a*), the KL divergence diverges to infinity (Equation 6). In practice, we add a small value *δ* = 10^−5^ to each distribution for all the state-action pairs of the environment the agent knows, and normalize each distribution so that the KL divergence is well defined. The KL divergence of the current model is always well-defined because the support of model *i*′ contains the latest *h*_*i*_*′* observations.

Based on the computation of KL divergences, there might be three outcomes, summed up in Equation 7. (1) If a known model explains the last observations better than the current model, the agent should replace its current model with a stored one. (2) If the current model does not explain the recent observations well, but is better than all stored models, the agent should create a new model. (3) Finally, if the current model has the minimal KL divergence and the divergence is low enough, the agent should keep using its current model. With 1 ≤ *i*′ ≤ *k* the current model, and the model creation threshold Δ_*c*_,

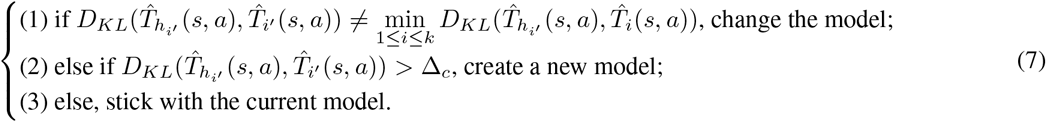

KL divergence helps detect that a change happened but does not indicate precisely when the change occurred. We explain how the agent can overcome this limitation in the next subsection which focuses on change point detection.

### 2.4 Retrospectively finding the change point

#### 2.4.1 Why should the agent find a change point?

Choosing the new model or context based on all recent observations is sub-optimal as it tends to favor moderate models. To illustrate this phenomenon, we take the example of a passive observer modeling a coin that can land on heads (*H*) or tails (*T*). The observer sees the sequence {*H, H, H, H, T, T, T*}, with tails being the latest observation. The agent has three coin models: Model 1, which is its current model and mostly predicts heads, Model 2, which mostly predicts tails, and Model 3, which predicts heads or tails with the same probability. After the last observation, Model 3 becomes the model which explains best the observed sequence and the agent must change its current model. Nevertheless, the agent should not use model 3 directly, although it explains all recent observations best. Instead, the agent must detect that the swap occurred between Model 1 and Model 2 after the last heads and before the first tails. Retrospectively finding this change point allows the agent to choose the appropriate associated model for the task.

A second implication of precise change point detection is that the agent can reassign observations to the correct models depending on when the change happened. Observations occurring after the change point belong to the new model and should be reassigned from the current model to the new model. This improves both models: the former becomes more precise by discarding incorrectly assigned observations, while the latter benefits from additional relevant experiences.

#### 2.4.2 Implementation of change point detection

To find the change point between the current model and a new one, the agent assesses to what extent it can sequentially redirect the latest observations from its current model to the new model (Figure 3). To find the change point, the agent gradually computes a cumulative sum of log-likelihood of observing recent observations for the current model and each potential new model. The agent must compute the likelihood of all possible change points and possible models. Hence, based on the number of stored models, this approach may possibly be computationally expensive.

**Figure 3:**
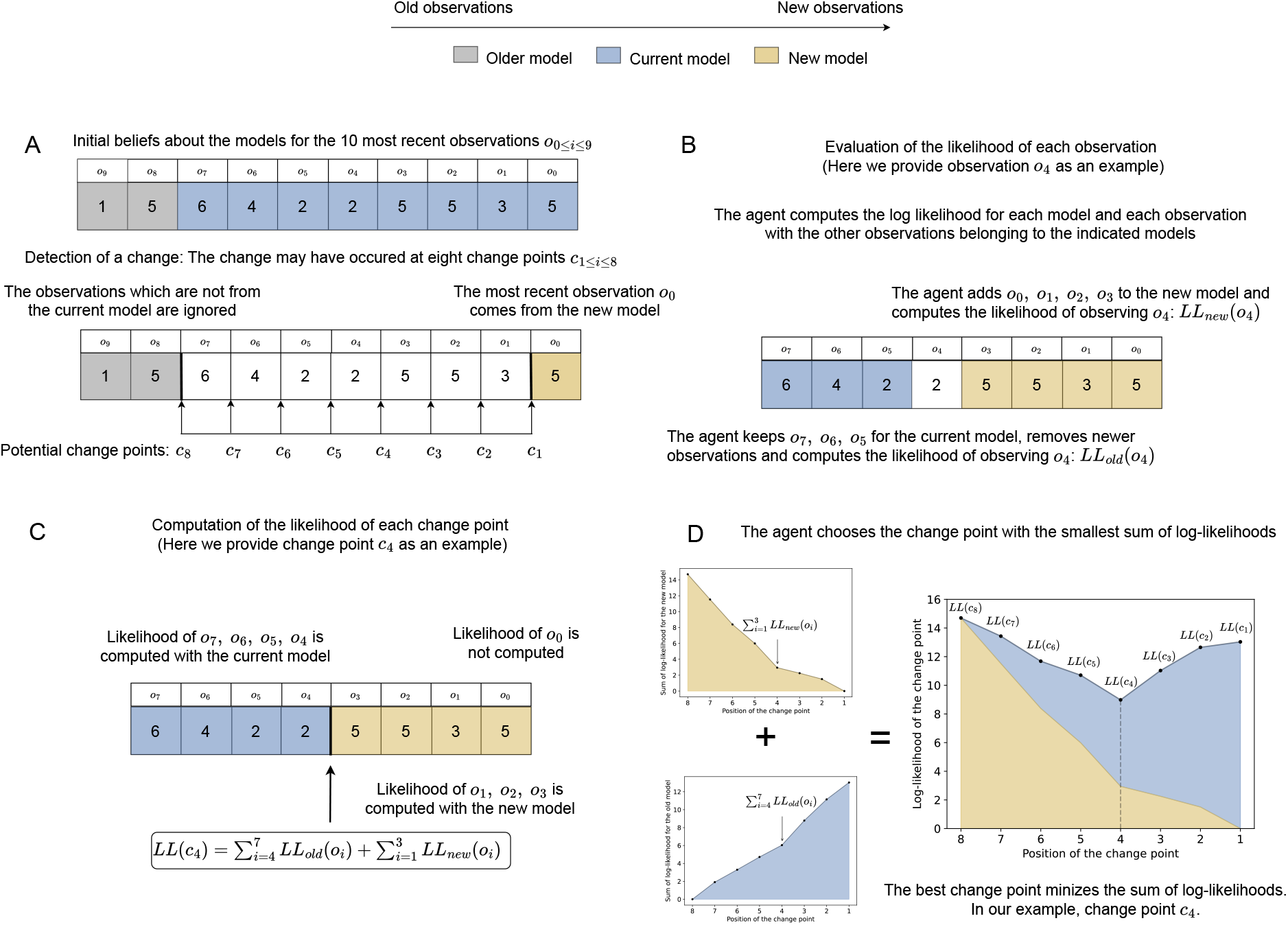
Illustration of the method to find a change point. The two distributions we compare are the top ones from Figure 4. When detecting a change, the agent follows these four steps (A to D) in order. **Panel A** The horizon has a size of *h* = 10, but the horizon for the current model alone is *h*_*i*′_ = 8 because an older model is used for the two first observations. As a result, the change point can only be at one of 8 possible positions. **Panel B** The agent computes the likelihood of each observation, for the two models. **Panel C** The total evaluation of each change point is the sum of the individual log-likelihoods the agent computed individually. **Panel D** The agent compares the log-likelihood of all change points. The change point which minimizes the sum of log-likelihoods in this example is change point 4.

We denote 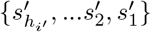 as the most recent arrival states the agent has observed from performing action *a* in *s*. Here, 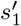 is the most recent observation while 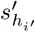 is the oldest. There are *h*_*i*_*′* 1 possible change points between two successive observations and two possible change points before or after all the observations. Nevertheless, the last observation belongs to the new model. Hence, there are *h*_*i*_*′* possible change points, going from only the last observation belonging to the new model to all of the last *h*_*i*_*′* observations belonging to the new model (Figure 3, Panel A).

To compute the log-likelihoods, the agent needs to be able to add the observations to the new model and to remove them from its current model. When the agent adds a new observation 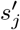 to a model *i*, the agent updates the transition function using Equation 8.

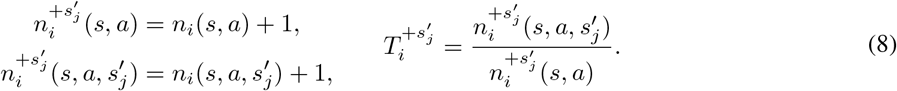

To remove an observation 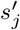 from a model *i*, all the pluses from Equation 8 become minuses. We extend Equation 8 to any ensemble of arrival states *A*(*s, a*). We denote 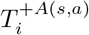 as the transition function of model *i* with the arrival states *A*(*s, a*) added to model *i*. Similarly, we denote 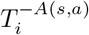 as the transition function when the arrival states *A*(*s, a*) are taken away from model *i*. Removing an observation from a model can only happen if the model has this observation in its model.

To compute how well the current model *i*′ explains the recent observations, the agent computes the log-likelihood of witnessing each observation supposing that the model used before the observation was *i*′ (Figure 3, Panel B). Starting with the second most recent arrival state 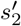 the agent computes its new transitions without 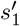 and 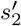 and computes the log-likelihood of observing 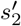 with model *i*′. Similarly, the agent computes the likelihood of observing each arrival state 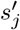 with 2 ≥ *j* ≥ *h*_*i*_*′* after removing 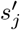 from its model and all the following observations. With 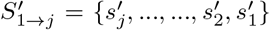 the ensemble of the last *j* arrival states, the likelihood of observing 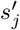 with the current model *i*′ is

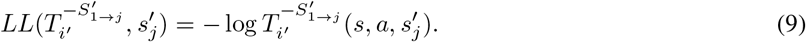

With this method, the agent can retrospectively compute the likelihood of observing each of the last *h*_*i*_*′* events with the model it had at the time of the observation. The agent does a similar computation for all the possible new models by adding the observations instead of taking them away from the model. Starting with the most recent observation 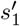, the agent computes the probability of observing 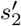. Similar to Equation 9, the likelihood of observing each event 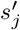 with the model *i* ≠ *i*′ is

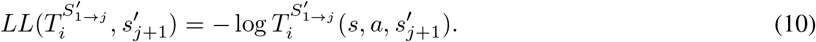

In Equation 9 and Equation 10, the agent adds *δ* = 10^−5^ to each distribution and normalizes them before computing the log-likelihood. This ensures that all the log-likelihoods are well defined. From Equation 9 and Equation 10, the agent can compute the likelihood of each change point. The likelihood of a change point *c* between model *i*′ and model *i* is given by the likelihood that observations before *c* belong to the current model *i*′, while those after *c* belong to the new model *i* (Figure 3, Panel C). For each couple of models (*i*′, *i*), the likelihood *L*_*i*_(*c*) of change point *c* with 1 ≤ *c* ≤ *h*_*i*_*′* reads

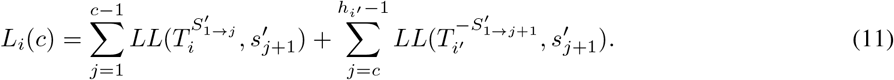

The optimal change point *c*_*opt*_ is defined as

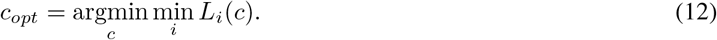

This method provides the new model *i*_*opt*_ and the optimal change point *c*_*opt*_ (Figure 3, Panel D). Once the agent finds the best change point and the best corresponding model, it reassigns the observations after the change point from its current model to the new model and starts using it (Figure 4).

**Figure 4:**
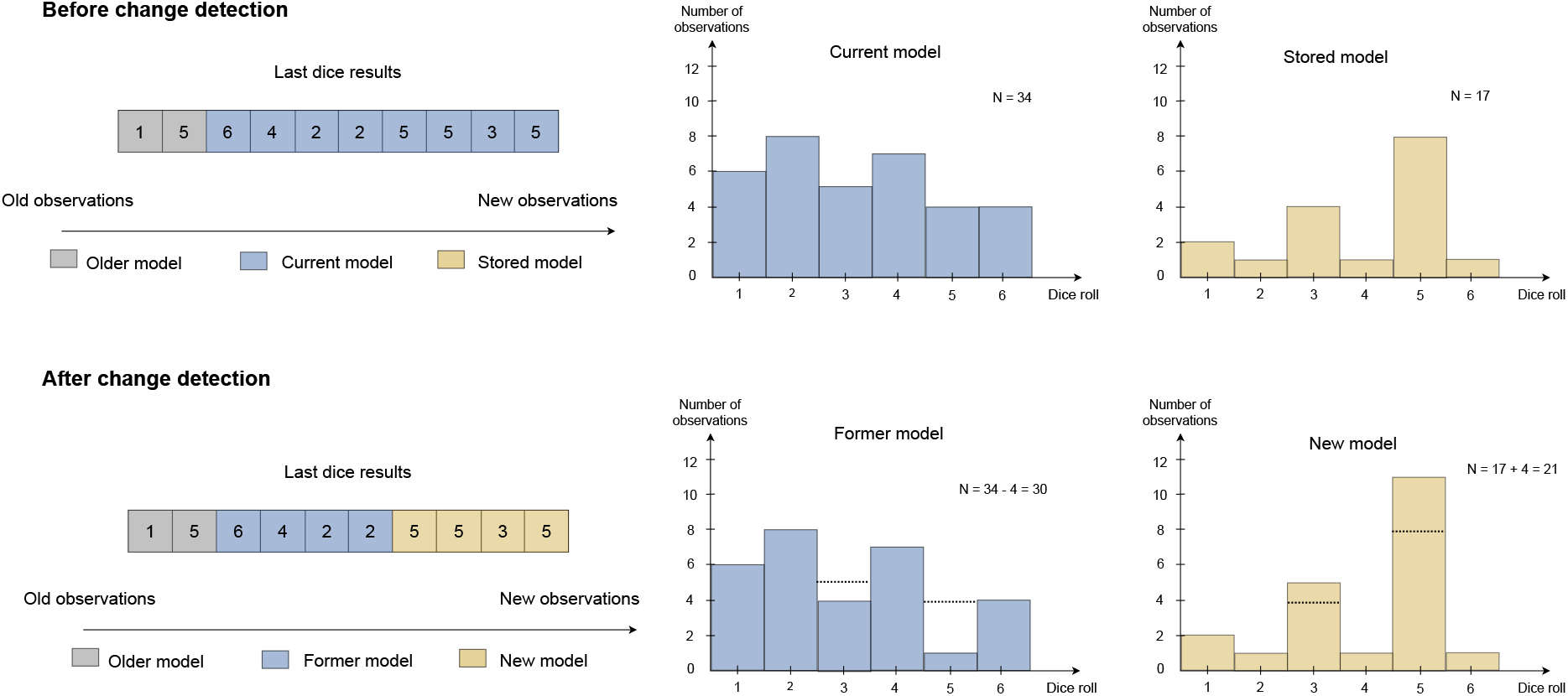
Illustration of the reassignment of the observations after finding a change point. (Left) The change point and the last observations come from Figure 3. (Top center and right) Histograms of the die rolls before the change detection for the current model and a stored model. We make the assumption that the agent detects a change from the current model to the stored model. (Bottom center and right) Histograms after the change detection. The current model becomes the former model and the stored model becomes the new model. The histograms illustrate the reassignment of the observations from the former model to the new model.

### 2.5 Merging similar models

#### 2.5.1 Why should the agent merge similar models?

There are several potential advantages to merge similar models. (1) Merging similar models reduces computational and memory costs, notably by limiting the number of comparisons when trying to detect a change or find a change point. (2) Merging models allows to have a more sensitive threshold for model creation, as similar models will be merged a posteriori. This may reduce the number of false negative errors, when the agent does not detect an existing change. (3) The capacity to merge models instead of forgetting them adds stability to MMRCPD, when the lack of stability is a common limitation of online and incremental methods [Beaumont et al., 2025].

Before merging two models, MMRCPD compares them to their combined distribution: if the combined distribution explains well both initial distributions, they can be merged. To do so, the agent uses a Jensen-Shannnon (JS) divergence, which is a common symmetric distance to compare distributions to their average distribution. The Jensen-Shannon divergence computes the KL divergences of the two distributions to their average distribution and sums them with an equal factor of 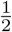. This distance has an upper bound of log_*b*_(2), where *b* is the base of the logarithm used in the KL divergence computation [Lin, 2002]. The existence of an upper bound on the distance facilitates model fitting and interpretability of the merging module. We present other potential merging implementations in the supplementary information (section A.1), such as weighted and semi-weighted JS divergences.

#### 2.5.2 Implementation of model merging

After every action *a* in state *s*, the agent checks if two models of (*s, a*) can be merged. If so, the agent merges the two models and continues merging models until no model can be merged anymore. When the agent checks whether it can merge two models or not, it computes a JS divergence between each couple of probability distributions. If the smallest JS divergence is below a merging threshold Δ_*m*_, the agent merges the two corresponding distributions. To compute the JS divergence between two transition distributions *T*_*i*_(*s, a*) and *T*_*j*_(*s, a*), the agent first computes the average distribution 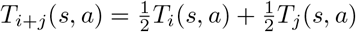. The JS divergence reads as

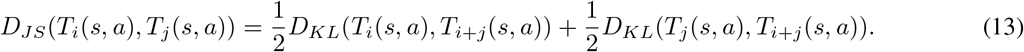

When the agent merges two models, it replaces one model with the merged model and forgets the other one, reducing the total number of models by one. The agent reallocates all observations from the discarded model to the merged model (Figure 5).

**Figure 5:**
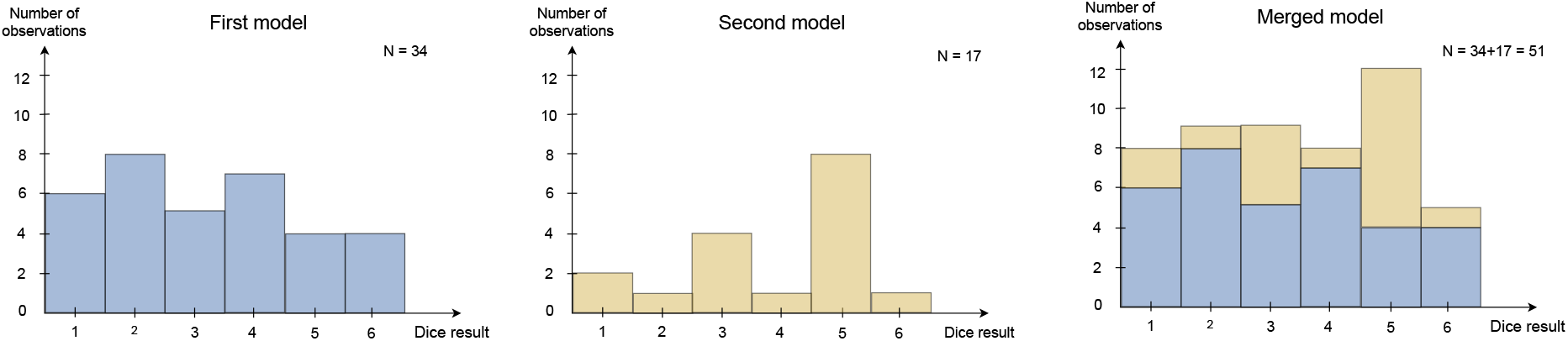
Example of the model merging process. If the initial models are similar to the merged model, the agent merges the two models by creating a model that includes all the observations. One can compute the probability distribution for each plot by normalizing the data by the the number of observations *N* .

### 2.6 Forgetting unused models

Forgetting models allows to limit the maximum number of models max_*mod*_. Forgetting models reduces memory and computational costs and aligns with biological constraints. For example, humans have limited working memory and may only maintain and evaluate a few strategies or models simultaneously [Collins and Koechlin, 2012].

MMRCPD forgets a model only when it stores its maximal amount of model max_*mod*_ and cannot merge any models. In this case, it discards the model with the fewest observations. If the current model is the smallest, the agent instead forgets the second smallest model. Retaining the current model ensures stability and reflects the idea that recent observations are more salient and more useful to solve the current task than older ones.

We chose to implement this forgetting mechanism for its simplicity. However, since the agent cannot forget its current model, it must be able to retain two local models per state-action pair and distribution type (transition or reward). In our simulations, the threshold max_*mod*_ depends on the state-action pair and the agent can store the same number of models per state-action pair. This forgetting module implies that max_*mod*_ ≥ 2.

There are at least two alternative implementations for the forgetting module. First, the threshold max_*mod*_ could become global rather than state-action specific. This would allow the agent to store more models in areas with frequent changes. Second, instead of forgetting the smallest model, the agent could forget the least recently used one, as models which have not been used for a long time may have become irrelevant.

## 3 Results

### 3.1 Environments

To simulate MMRCPD, we mainly use three classes of environments (Figure 6). The first one consists in small one-step environments with three or four states and two actions. The second one is a chain task, with 5 states and 2 actions. Finally, the third type of environment is an ensemble of 7 × 7 maze environments with random wall generation. For each environment, we simulate volatile and uncertain variations.

**Figure 6:**
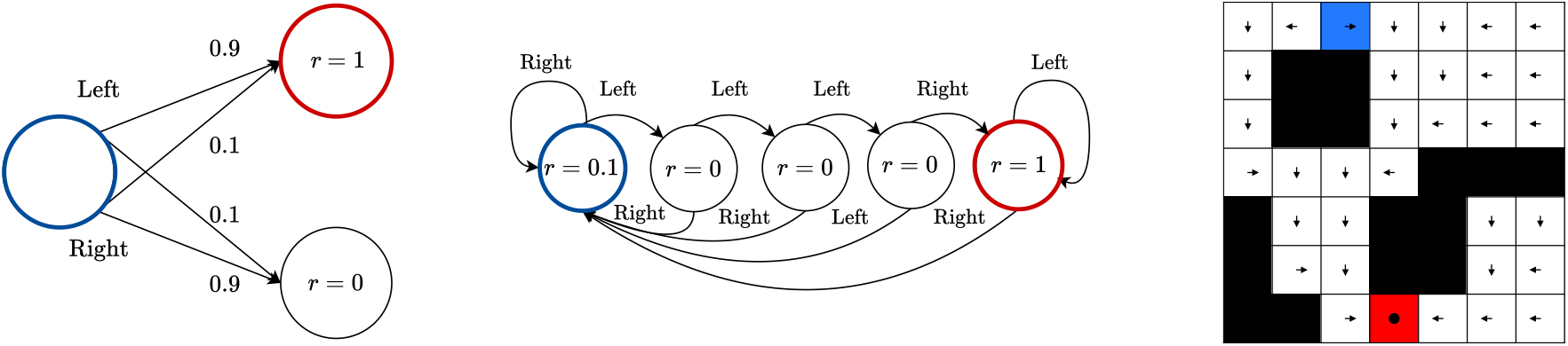
The three types of environments used in this article. In all the environments, the initial state is highlighted in blue and the most rewarded state in red. The agent should learn to go from the blue initial state to the red one and exploit it. (Left) The one-step environment. The agent must choose between two actions, *Left* or *Right*. The two actions have a slipping probability *p*_*slip*_ = 0.1 to result in the opposite action. (Center) The chain task, which is a 5-state and multi-step extension of the one-step environment. The slipping probabilities (*p*_*slip*_ = 0.1) of the *Left* and *Right* actions are not represented for readability. (Right) One of the 10 mazes the agent faces of size 7 × 7, with randomly generated walls, in black. The agent can perform one-step movements in the four cardinal directions, and a *Stay* action to remain in its current cell. One-step movements are indicated with arrows and the *Stay* action with a filled circle. The actions represented correspond to the optimal policy to reach the rewarded state and exploit it.

#### 3.1.1 The one-step environments

##### Base environment

The base version of the one-step environment is a three states (one initial state and two arrival states) and two actions (*Left* or *Right*) environment (Figure 6, Left). The goal of the agent is to reach the state with a positive reward *r >* 0 by choosing the *Left* action that leads to it. **Base stochasticity:** For all experiments, the agent has a probability of slipping *p*_*slip*_ = 0.1, which causes it to experience the outcome (reward and arrival state) of the action it did not choose. After one step, the agent is reset to the initial state. **Volatile variation:** The *Left* and *Right* actions are swapped at a fixed rate (Figure 7, Center-Left). **Uncertain variation:** This variation consists in adding a third arrival state the agent reaches 10% of the time when choosing the correct action. Agents can only get a reward of *r* = 10 when reaching this state, meaning that they must learn an infrequent event to find the optimal policy (Figure 7, Top-Left).

**Figure 7:**
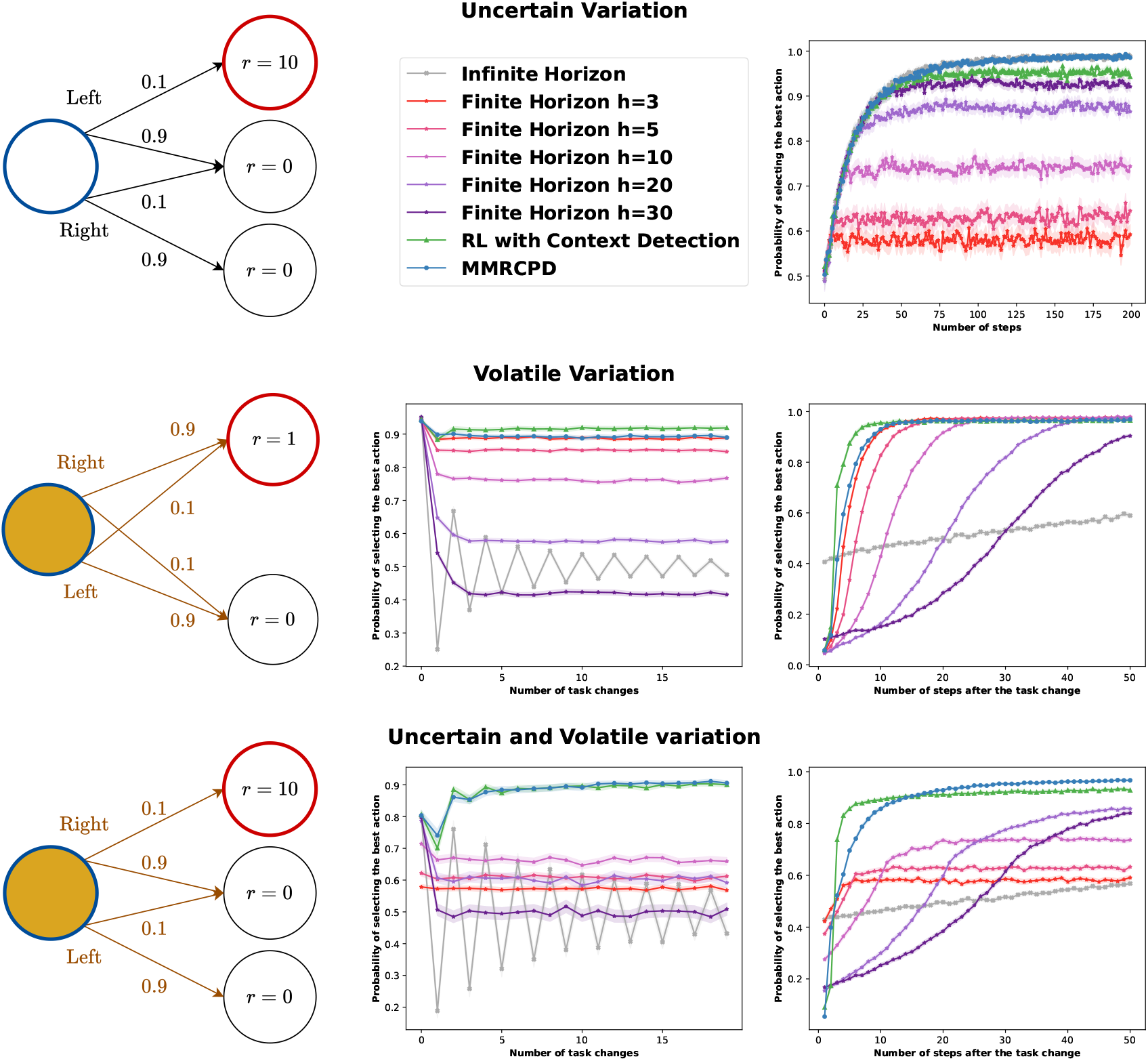
Comparison of performance of the agents in the uncertain (Top), the volatile (Center), and the uncertain and volatile (Bottom) variations of the one-step environment. (Left) Sketch of the environment used. (Top-Right) Probability of choosing the best action in the uncertain variation of the task. In this experiment, MMRCPD (blue) and the infinite horizon agent (grey) overlap. For volatile variations, we show two plots: the evolution of the averaged probability of choosing the best action over time (Center) and after a task change (Left).

#### 3.1.2 The chain environment

##### Base environment

The chain task [Dearden et al., 1998, Poupart et al., 2006] is an environment of intermediate complexity, where the goal of the learning agent is to reach the farthest state from its starting state to get a maximal reward of *r* = 1 (Figure 6, Center). Every time the agent chooses the action *Left*, it goes back to the initial state where it receives a bait reward of 0.1. If the agent reaches the farthest state from its starting position, it receives a reward of *r* = 1. The optimal policy is to do the action *Right* only. The chain environment we implemented has 5 states and 2 actions. **Base stochasticity:** For all experiments, the agent has a probability of slipping *p*_*slip*_ = 0.1 and receiving the outcome (arrival state and thus reward) of the opposite action. **Local volatility:** The task-change we introduce is a local task change, with the *Right* and *Left* action inverted for one state every 10 trials of 50 steps (Figure 8, Top-Left). Thus, there are 2^5^ = 32 possible combinations or contexts. **Global volatility:** The *Right* and *Left* action are inverted everywhere in the environment (Figure 8, Center-Left). **All-but-one volatility:** The *Right* and *Left* action are inverted everywhere in the environment but for one state randomly sampled at every change (Figure 8, Bottom-Left).

**Figure 8:**
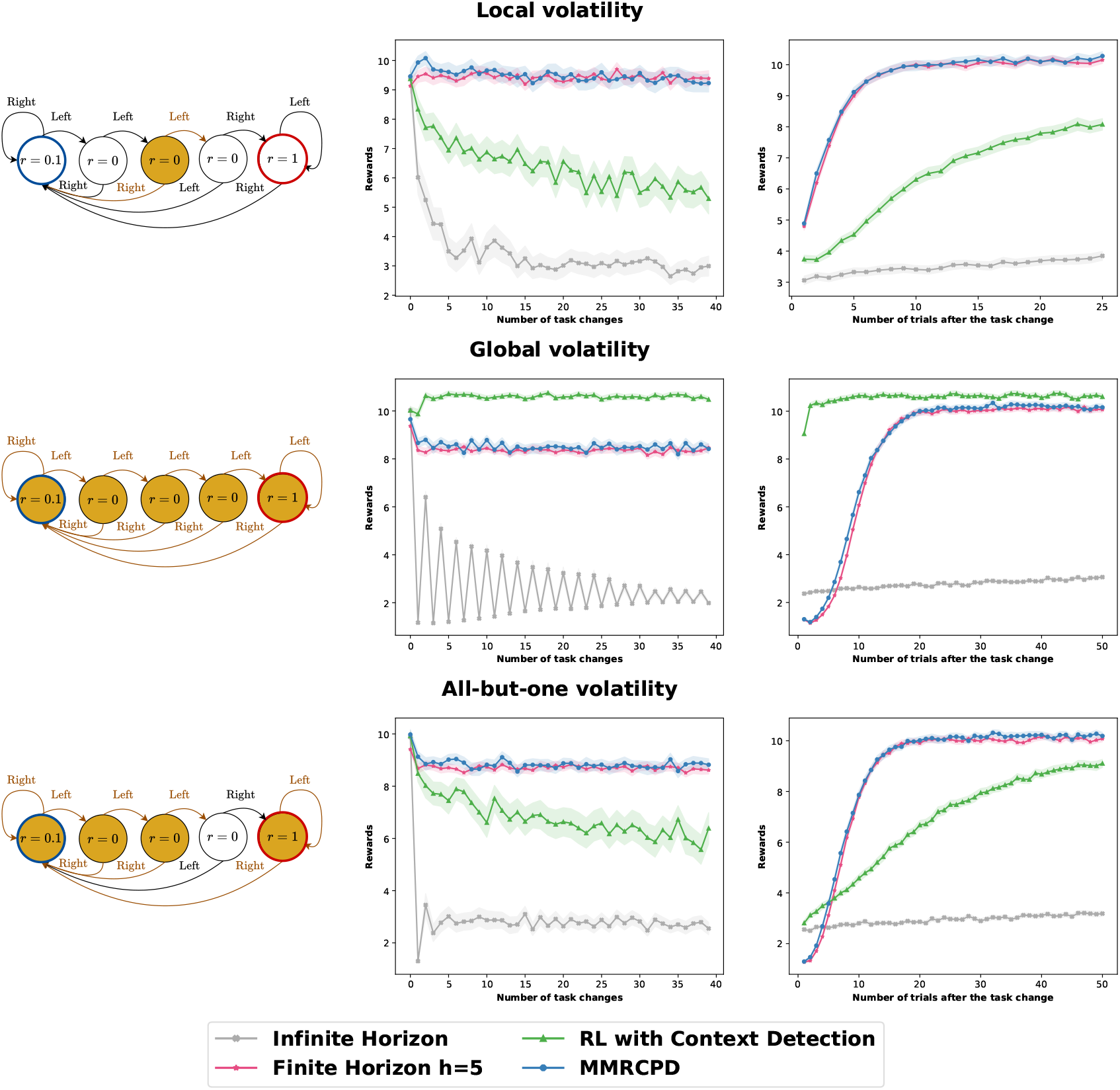
Comparison of performance of local- and context-level approaches based on the nature of the change. (Left) Sketch of the environments. (Center and Right) Averaged results over time and after a task change. (Top) When the change is local, the context-level method fails to adapt to volatile environments. (Center) When the change is contextual, the context-level method outperforms the local change detection methods, as it detects the task change faster. (Bottom) If the change is global but for one randomly selected state, context-level methods fail to adapt to changes.

#### 3.1.3 The maze environments

##### Base environments

The maze environments are 7 × 7 squared environments with random wall generation (Figure 6, Right). The goal of the agent is to reach the maximal reward of 1 and to exploit it. The cell with the reward is 10 cells away from the initial cell. The agent can perform 4 one-step movement actions (*Up, Down, Left, Right*) or the *Stay* action. **Base stochasticity:** For all experiments, the transitions are slightly stochastic. When the agent performs one of the 4 one-step movement actions, it has a 75% chance of reaching the state corresponding to the deterministic action and a 5% chance of ending in nearby cells (Figure S2). When the arrival state of the agent is a wall, the agent stays in its starting state. The *Stay* action is deterministic with a 100% chance of staying in the same cell. We provide all the simulated mazes in the supplementary information (Figure S1). **Volatile variation:** For each task change, the effects of all the actions in 20% of the cells are changed, following a predefined cyclic permutation for each cell (Figure 9, Center-Left). There are two possible distributions per cell (either the classical one or a cyclic permutation of it). Thus, there are theoretically 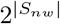 possible contexts, with |*S*_*nw*_ | the non-wall states of the maze. The example maze (Figure 6, Right and Figure 9, Left) has 2^39^ possible contexts. Therefore, a context-level agent cannot learn a model of the environment for all of the different contexts. **Uncertain variation:** For all actions, agents have a 5% chance of being reset back to their starting position (Figure 9, Top-Left). Agents must learn to ignore this uncertain event.

**Figure 9:**
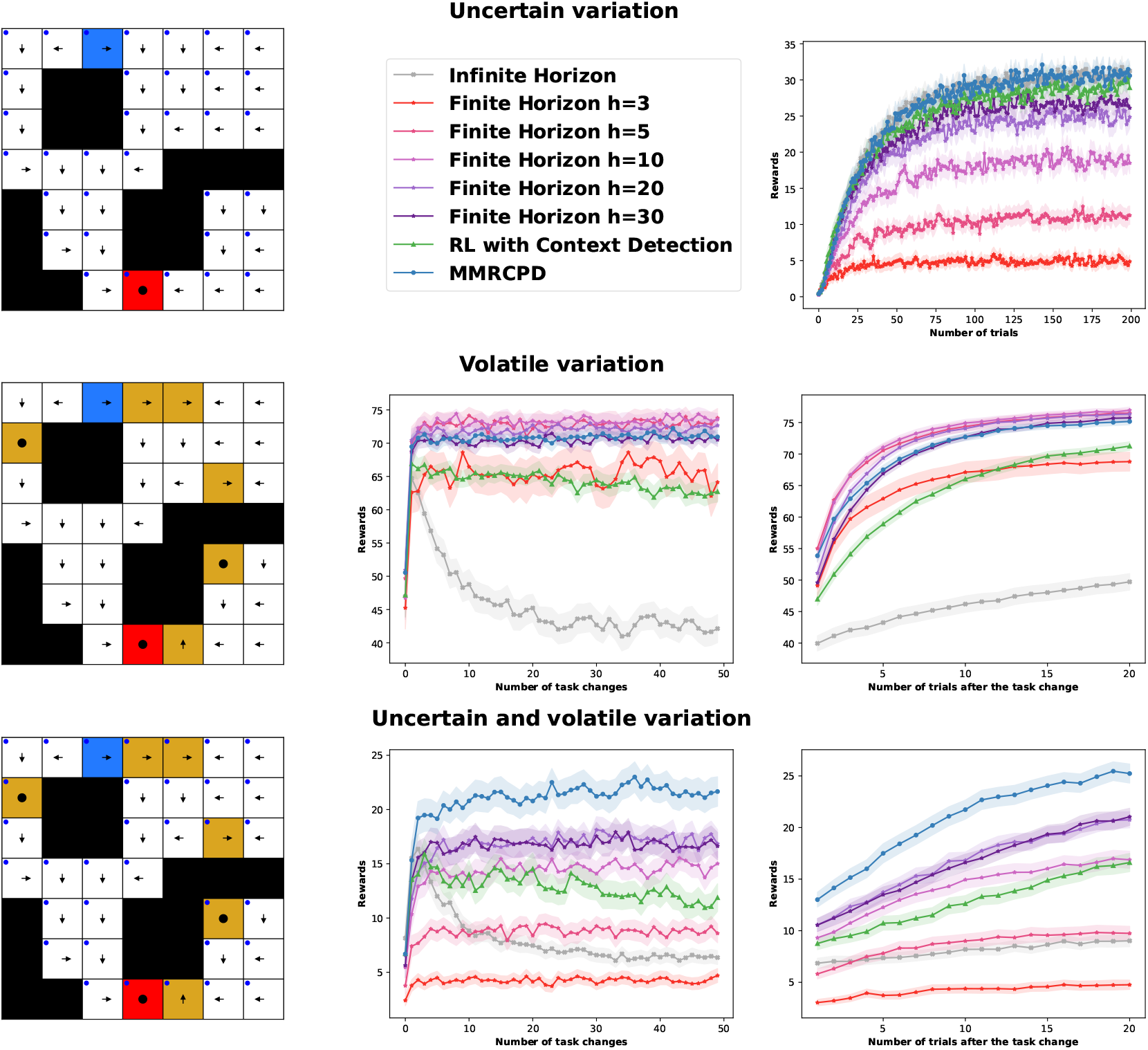
Comparison of performance of the agents in the uncertain (Top), the volatile (Center), and the uncertain and volatile (Bottom) variations of the mazes. (Left) Sketch of the environments. (Top-Right) Performance of the agents in the uncertain mazes. For volatile variations, we show two plots: the average performance over time (Center) and the average performance after a change (Right).

### 3.2 Simulations

We compare MMRCPD with infinite-horizon (Equations 1 and 2) and finite-horizon agents (Equations 3 and 4), both using a single model per state-action pair. We also compare MMRCPD to a context-level change detection agent, named RLCD, which stands for RL with Context Detection [Da Silva et al., 2006]. All agents are model-based and use the same softmax decision-making process for fair comparison. All simulations show the mean performance and the confidence interval at 95%.

MMRCPD can theoretically adapt to changes at any level, from local variations to broader context shifts, provided they do not happen too frequently. For example, if one local change happens faster than the study horizon *h*, MMRCPD cannot detect this local change and averages out changes. To simplify our plots and analyses, we use environments where dynamics change at a fixed rate. No agent knows nor explicitly uses this information.

The number of states and actions in a chosen environment remains constant over task changes. All of these changes pertain specifically to transitions. Hence, MMRCPD and RLCD maintain a reward model for each transition model and do not try to detect changes in reward. When they change their transition models, they change their reward models accordingly. Notably, MMRCPD reassigns the rewards to the appropriate model based on the detected change point.

### 3.3 Adaptation to uncertain and volatile tasks

To illustrate the advantage of change detection methods over finite- and infinite-horizon agents, we simulate the agents in the one-step environments in the uncertain, volatile, and uncertain and volatile conditions. For the uncertain variation, we examine the performance of the agents over 200 trials and 2000 seeds. For the volatile and uncertain-volatile variations, we swap actions *Left* and *Right* every 50 steps over 2000 trials, on 500 seeds. All agents have the same exploration rate *β* = 10. The finite-horizon agents use an horizon of *h* = 3, *h* = 5, *h* = 10, *h* = 20, or *h* = 30. RLCD uses *h* = 50, *Emin* = − 0.05, *ρ* = 0.4, and Ω = 0. MMRCPD uses *h* = 3, Δ_*c*_ = 0.5, Δ_*m*_ = 0.3, and max_*mod*_ = 5. We keep this set of parameters for all the simulations in the one-step environments.

In the uncertain variation of the one-step environment (Figure 7, Top), short-horizon agents do not perform well as they tend to forget the infrequent, yet key transition to the rewarded state. The longer the horizon, the better finite-horizon agents perform. Consequently, the infinite-horizon agent, which averages out the uncertainty of the environment, finds the optimal policy. MMRCPD performs as well as the infinite-horizon agent. This illustrates that adding the capacity to detect task changes and maintaining multiple models does not impair the performance of MMRCPD in stable environments. RLCD is slightly less stable than MMRCPD and the infinite-horizon agent, but performs almost as well.

The finite-horizon agents with short horizon, RLCD and MMRCPD adapt well to task changes (Figure 7, Center). Long-horizons agents adapt poorly to task changes, and the infinite-horizon agent averages out all the changes. Hence, the long horizon agents which performed best in the uncertain variation of the task perform worse in its volatile variation. RLCD performs best in this volatile variation, as learning a context facilitates the change detection on the *Left* action when the *Right* one changes, and vice versa.

When combining both variations, the finite- and infinite-horizon agents fail to adapt to the uncertainty and the volatility of the environment respectively (Figure 7, Bottom). Therefore, all single-model agents fail to select the optimal action and end up with a poor performance. MMRCPD and RLCD, which maintain accurate models of the environment while being able to detect task changes, perform well when combining volatility and uncertainty. Importantly, their performances increase over time, indicating that their models of the environment improve. RLCD detects changes faster than MMRCPD, while being slightly less stable. As actions *Left* and *Right* change at the same time, RLCD can maintain two contexts only and detect contextual changes faster. However, RLCD and similar context-level agents cannot perform well when the changes are local and not contextual.

### 3.4 Contextual vs. local volatility

To compare how contextual and local change detection adapt to various volatile environments, we simulate RLCD and MMRCPD in the chain environments over 100 seeds. Local changes happen every 500 steps, while the global and all-but-one changes happen every 1000 steps. The task is episodic and the agents are put back to their initial state after 20 steps. RLCD uses *h* = 20, *Emin* = − 0.05, *ρ* = 0.3, and Ω = 0. MMRCPD uses *h* = 5, Δ_*c*_ = 1, Δ_*m*_ = 0.1, and max_*mod*_ = 5. We keep this set of parameters for all the simulations in chain environments. We also use an infinite-horizon and a finite-horizon agent with *h* = 5 as baselines. All agents use *β* = 3.

RLCD fails to adapt to locally changing environments (Figure 8, Top). The performance of RLCD worsens over time, as it cannot make sense of local changes and ends up averaging out changes. MMRCPD and finite-horizon agents adapt well to locally changing environments. When facing a two-context environment (Figure 8, Center), RLCD learns the two contexts well, and outperforms other agents. However, MMRCPD and finite-horizon agents still manage to learn a close to optimal policy, and their performance do not decrease over time: they manage to learn the different changes, but are slower at detecting them. In the discussion, we propose several ways to fasten context change detection for MMRCPD, *e.g*., by associating local changes together. Interestingly, if all states change but one random state, RLCD already fails to adapt to the changes (Figure 8, Bottom). MMRCPD and finite-horizon agents manage to learn a good policy, at a slower pace than when the change is local. These results illustrate that context-level methods do not adapt well to locally changing environments, whether the change concerns one randomly selected state, or almost all of them.

### 3.5 Adapting to large and locally-changing environments

To further demonstrate the interest of our method, we compare MMRCPD to other agents in large, uncertain, and locally changing mazes. We evaluate the agents in 10 mazes over 50 seeds for the uncertain variation, and 20 seeds for the volatile and volatile-uncertain variations. All agents use *β* = 3 and the multi-model agent uses *h* = 10, Δ_*c*_ = 1, Δ_*m*_ = 0.1, and max_*mod*_ = 5. We test finite-horizon agents with *h* = 3, *h* = 5, *h* = 10, *h* = 20, and *h* = 30. RLCD uses *ρ* = 0.2, *h* = 50, *Emin* = − 0.05, and Ω = 0. One trial lasts 100 steps, and the environment changes every 2000 steps in the volatile variations.

Short-horizon agents do not adapt well to rare events in the mazes (Figure 9, Top). Acting in a large environment does not impair the performance of MMRCPD, which acts as well as the infinite-horizon agent when the environment is stable. RLCD is slightly less stable than these two agents. The results in the uncertain variation of the mazes echo the ones presented in the one-step environments (Figure 7).

MMRCPD and finite-horizon methods adapt well to local changes in mazes (Figure 9, Center). Finite-horizon methods with very short or very large horizon perform slightly worse than methods using a rolling window of intermediate size, which may come from the stochasticity of mazes even in the volatile variations (Figure S2). MMRCPD performs almost as well, yet slightly worse than some finite-horizon agents with *h* = 5, *h* = 10, or *h* = 20. RLCD and the infinite-horizon do not adapt well to local changes.

When combining both variations, MMRCPD clearly outperforms all other agents (Figure 9, Bottom). In particular, MMRCPD is the only agent whose performance increases over time: finite-horizon agents quickly reach a plateau, while RLCD and the infinite-horizon agent worsen over time. This further highlights that MMRCPD is a versatile agent, which can adapt to uncertain, volatile, and uncertain and volatile environments. Combined with the simulations in the one-step environment and the chain task, these results show that MMRCPD adapts well to environments of various sizes, with various uncertainty and volatility types.

### 3.6 The interest of retrospective change point detection

Retrospective change point detection helps the agent maintain accurate models of the task by reassigning the observations to the correct models. This property of MMRCPD seems particularly useful when the task changes fast, or when the environment is uncertain and the agent needs several observations to detect a change. The interest of retrospective change detection grows as the number of changes increases: without precisely finding change points, learned models may spill on each other. To illustrate the interest of retrospective change point detection, we simulate an ablation of MMRCPD which only reassigns the last observation to the new model when detecting a change, and compare it to MMRCPD on the chain task with local volatility. We increase the level of volatility on the chain task by reducing the number of steps between two changes from 500 to 100. We do not change the parameters of MMRCPD used previously in the chain task (Figure 8, *h* = 5, Δ_*c*_ = 1, Δ_*m*_ = 0.1, max_*mod*_ = 5). The ablation of MMRCPD which does not reassign the observations to the correct models after finding the change point performs worse than MMRCPD (Figure 10). This performance decreases over time as models deteriorate.

**Figure 10:**
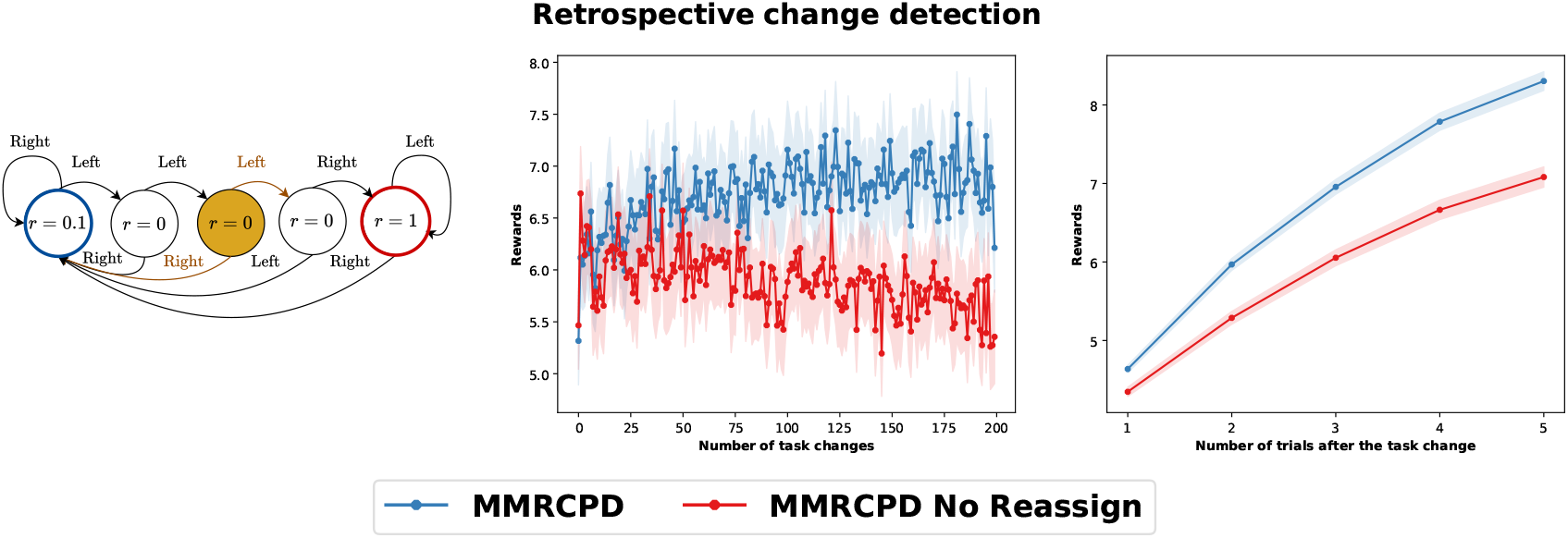
Illustration of the interest of the retrospective change point finding. MMRCPD and an ablation of MMRCPD, which only reassigns the most recent observation to the new model when changing models, are compared on the locally changing chain task. (Left) Sketch of the environment. (Center) Averaged performance of the agents over time. (Right) Averaged performance of the agents after a change.

## 4 Discussion

### 4.1 Conclusions

We introduced a novel multi-local-model approach to deal with local task-changes at a lower memory cost than context-based approaches (Figure 1). Based on this local-model-level approach, we presented a model-based reinforcement learning agent, named Multi-Model with Retrospective Change Point Detection (MMRCPD), which arbitrates between multiple local models (Figure 2). This agent can create, merge, and forget models at the level of the state-action pair based on the environment it interacts with. Additionally, it retrospectively finds change points after change detection to reassign past observations to the correct models (Figures 3, 4 and 5). We compared MMRCPD with context-based, infinite-horizon, and finite-horizon agents in volatile and uncertain environments of various types and sizes (Figures 6). MMRCPD outperforms other agents in locally changing and uncertain uncertain environments, and performs well in all environments (Figures 7, 8, 9). In the supplementary information, we provide extended methods, show that the parameters of MMRCPD are explicable and stable, and demonstrate that MMRCPD has limited computational cost.

### 4.2 Comparing MMRCPD with existing methods

Inferring contexts and adapting to them is a candidate to explain human behavior in many sub-fields of cognitive sciences, such as classical conditioning, episodic memory, instrumental learning, and motor adaptation [Heald et al., 2023]. In our work, we focus on decision-making and reinforcement learning, with an agent acting in a Markov decision process and trying to infer the current structure of the task to solve it. A common way to detect changes in the decision-making literature is to use a Bayesian inference approach, *e.g*., using Dirichlet processes or Kalman filters [Gershman et al., 2010, Collins and Koechlin, 2012, Heald et al., 2021, Velentzas et al., 2017]. We compare the characteristics of our approach to two Bayesian inference approaches: the PROBE [Collins and Koechlin, 2012] and the COIN methods [Heald et al., 2021].

MMRCPD does not share many features with COIN. For example, COIN updates all contextual models at the same time, using a weight based on their reliability (or responsibility in the original article). Contrarily, our method updates its current model only, and corrects the wrongly assigned observations with retrospective change point detection once the change is detected. COIN mainly focuses on motor learning, while our approach aims to model behavior in a decision-making task, similar to PROBE. The PROBE approach shares more features with our approach as it builds different task-sets, arbitrates between them online based on their reliability and discards new models that are outperformed by stored models. Nevertheless, there are several differences between the two methods. First, PROBE works at the context level, switching the current strategy based on the incoming observations, while our agent works at the local transition level using model-based reinforcement learning. Second, MMRCPD retrospectively detects the moment the switch happened, whereas PROBE only adds the latest observation to the new context. Third, we do not compare the reliability of the agent’s strategy to the reliability of random behavior. Instead, our agent uses a sliding window for the latest observations and checks if its current model explains them. Fourth, we use a merging module that can merge similar models, while there is no merging module in the PROBE approach. This merging module limits the number of models, which leads to little model discarding while the PROBE method often discards unreliable newly created strategies, reducing its stability [Beaumont et al., 2025]. Finally, our approach scales well to large environments and can be applied to all Markov decision processes. Inversely, Bayesian approaches often scale poorly to large environments and focus on smaller problems such as classical conditioning or multi-armed bandits [Velentzas et al., 2017, Gershman et al., 2010].

In the reinforcement learning literature, several algorithms tried to adapt to task changes with online arbitration between models. The Multiple Model-Based Reinforcement Learning (MMRL) agent [Doya et al., 2002] uses a responsibility signal to weight the outputs of multiple specialist modules. This responsibility signal changes based on how well each specialist predicts the next state. Contrary to our method, MMRL uses a fixed number of models. To solve this limitation, Reinforcement Learning with Context Detection (RLCD) arbitrates between different contexts without knowing the number of contexts in advance [Da Silva et al., 2006]. RLCD uses an aggregate measure of discrepancy between the stored reward and transitions and the current experience. RLCD has several parameters: *h* (*M* in the original article), the horizon for model learning which is also used in change detection; *Emin* (*λ* in the original article), the threshold for context creation; *ρ*, a learning rate to update the predictive error of each context; and Ω, to weight change detection in transitions vs change detection in rewards. When no stored context can explain recent observations, RLCD creates a new uniformly initialized context. As a result, the memory usage of RLCD increases linearly with the number of contexts. More recent methods have either tried to improve RLCD by modifying the change detection method [Hadoux et al., 2014], demonstrated theoretical properties of a worst-case scenario algorithm using tree search when the environment changes gradually [Lecarpentier and Rachelson, 2019], or focused on detecting changes with a model-free approach [Padakandla et al., 2020]. In model-based reinforcement learning, at least one Bayesian approach detects changes in transitions in a complex and continuous environment [Velentzas et al., 2023]. However, all of these methods use a context-level approach and not our local-model-level approach (Figure 1). Thus, although they may adapt faster to contextual changes as shown with RLCD (Figure 8, Center), they fail to adapt to the exponential number of contexts in the environments we introduced (Figure 8, Top and Bottom, or Figure 9).

### 4.3 Limitations and future directions

MMRCPD is a simple and stable multi-model approach which can be modified in various ways based on the problem to solve. Notably, since MMRCPD is modular, one can adapt the learning, the forgetting, the merging, the change detection, or the retrospective change point finding modules to their specific problem, without modifying the rest of the agent.

#### Detecting changes in rewards

There are two main differences between transition and reward change detection: rewards are one-dimensional and ordered while arrival states form an unordered multi-dimensional space. For transitions, the sampling approach we used provides a distribution (an histogram) over arrival states. This same approach for the reward function provides a point-estimate value only (the mean of the reward distribution). To apply directly MMRCPD to rewards, one can use a distributional approach to learn the reward distribution. Additionally, once can change some metrics we used to adapt them to ordered distributions. In the reinforcement learning literature, a few algorithms try to detect changes in reward [Da Silva et al., 2006, Hadoux et al., 2014]. These works often use prior information on the reward distribution to detect changes in rewards, such as the maximal and minimal value of the reward distribution [Da Silva et al., 2006], or all the potential discrete reward values [Hadoux et al., 2014]. If the agent tries to detect changes in rewards and in transitions simultaneously, it can combine change detection metrics to detect joint changes.

#### Adapting to drifting environments

In their everyday lives, animals sometimes face drifting environments with a small but steady change which becomes prominent over time. For example, the amount of water in a pond may decrease slightly over time based on weather conditions and the numbers of animals drinking out of it. MMRCPD does not take into account drifting situations as it focuses on detecting abrupt local changes. To consider drifting environments, MMRCPD could use a finite, long horizon (of size *H >> h*), instead of an infinite horizon for the models it stores. MMRCPD would remain similar to what was presented before, with this additional parameter to account for drifts.

#### Directing exploration based on model reliability

MMRCPD does not make any assumption on how the agent makes a decision. Hence, it can use exploration bonuses to modify its behavior. When creating a new model or when a model has little information, the agent may explore to gather information and add observations to the model. To implement this, the agent can directly use optimistic in the face of novelty methods which come from the count-based exploration literature for single-model agents [Brafman and Tennenholtz, 2002]. These methods bias the behavior of the agent to gather more information on models which have little observations. The multi-model agent can also use model reliability to favor exploration. With our approach, the KL divergence between the recent observations and the current model directly indicates the reliability of the current model. A decrease in the reliability of the current model may indicate that there is a surprising observation the agent needs to investigate. An important property of using model reliability as an incentive to explore is that this measure decreases when the model of the agent matches recent observations well. This avoids that the agent remains permanently motivated to explore something inherently uncertain, which is a common limitation in the intrinsic motivation literature [Gottlieb and Oudeyer, 2018, Chartouny et al., 2025].

#### Associating local changes for exploration and context change detection

MMRCPD retrospectively infers the precise time-points when changes happened. Hence, it can compute the temporal co-variation of local changes, for example using a co-variation matrix. Once the agent learns a co-variation matrix and detects a change for a given state-action pair, it can re-explore all of the state-action pairs which correlate with it. This may yield a faster change detection, by directing exploration towards areas which are known to vary together. This possibility to detect precisely when local changes happen and associate them also allows to form contexts with a bottom-up approach.

#### Explaining human behavior

Our approach applies to all Markov decision processes and could thus be used to describe human behavior in many decision-making tasks such as multi-armed bandits or two-step tasks. Notably, our method may apply well to tasks with partial changes, such as the monkey feeding task [Beaumont et al., 2025]. In this task, human participants face three monkeys which have fruit preferences and need to identify which fruit each monkey prefers. The experimenters gathered results with complete rule changes (all monkeys change their preferred fruit) and with partial rule changes (one or a few, but not all monkeys, change their fruit preference). Task changes are not indicated to the participants: they must infer the best strategy based on the monkeys’ observed preferences only. Our model may adapt well to such tasks with local changes, and provide theoretical predictions such that reaction time increases after a task change (Figure S7), or that humans adapt better to recurrent changes than new ones. Additionally, the experimenters observed that the performance of participants decreases while exploration increases after a partial change, even for stable fruit-monkey associations. This indicates that changing the environment somewhere modifies the behavior of the agent in non-modified parts of the environment. Adding co-variation exploration and inference after a detected local change to our model would provide predictions on when and where this effect may happen.

## Acknowledgments

This research was funded in whole, or in part, by the French Agence Nationale de la Recherche (ANR) (ANR-21-CE33-0019 ELSA project). We would like to thank Olivier Serris for discussions on the change point detection method, and Abhishek Banerjee, Oriane Demandolx, Clarissa Montgomery, Geoffroy Morlat, and Matthias Tsai for their comments on a previous version of the manuscript.

## Copyright

Article and figures by Chartouny, Khamassi, and Girard (2025); available under a CC-BY 4.0 license.

## Code

The code is available online on github.

## A Extended methods

### A.1 Merging distributions for MMRCPD

Although we used the Jensen-Shannon divergence for MMRCPD in the main article, we tested a few variations, such as weighted [Corander et al., 2022] and semi-weighted Jensen-Shannon divergences.

The Jensen-Shannon divergence of two probability distributions *P* and *Q* is defined as

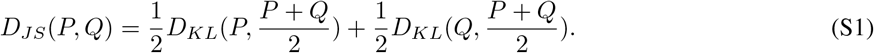

However, the Jensen-Shannon divergence in Equation S1 takes the average of the two probability distributions as a merged distribution. In our scenario, since we take the size of the distributions in the merging process (Figure 5), we may want the merging conditions to account for the relative sizes of the two distributions. Instead of comparing the two distributions to their average, we can compare them to a weighted merged distribution where each distribution’s influence is proportional to its number of observations.

A distance measure inspired by the Jensen-Shannon divergence, which weighs the initial distributions during merging is the weighted Jensen-Shannon divergence. This method maintains some theoretical guarantees of the Jensen-Shannon divergence, such as an upper bound on the divergence [Corander et al., 2022]. It also accounts for the size of the distributions in the divergence computation by replacing the 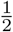 factors from Equation S1 with weighted factors that depend on the sizes of the two distributions. Given *N*_*P*_ and *N*_*Q*_ as the weights of distributions *P* and *Q* (*e.g*., their respective model sizes), the weighted Jensen-Shannon divergence is defined as

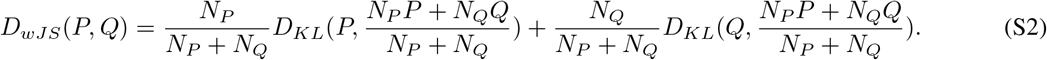

However, when one distribution has significantly more observations than another one, their weighted Jensen-Shannon divergence is close to 0 (Equation S2). In our scenario of model merging, weighting the KL divergences by the number of observations would lead to large models overpowering smaller ones, reducing model diversity. To address this, we introduce the semi-weighted Jensen-Shannon divergence, which uses weighted merging but keeps the KL divergence factors unweighted. The semi-weighted merging Jensen-Shannon divergence is

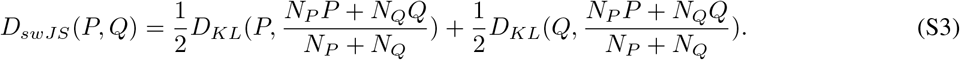

Using the semi-weighted merging JS divergence from Equation S3 instead of the classical JS divergence has several implications. For example, we lose some theoretical properties of the Jensen-Shannon divergence and its weighted alternative, such as upper bounds. However, this method may preserve model diversity better than JS, as it incorporates model sizes into the merging process. If one model *P* is much bigger than one model *Q*, the semi-weighted Jensen-Shannon divergence is close to a KL divergence (with a factor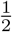). Unlike the KL divergence, the semi-weighted Jensen-Shannon divergence is always well-defined here because the merged distribution includes the observations from both distributions. If the two distributions are of the same size, the semi-weighted Jensen-Shannon divergence reduces to the Jensen-Shannon divergence. In our simulations, we did not observe that semi-weighted JS performed better than JS, and chose to stick to the JS divergence for its theoretical properties.

### A.2 Mazes

The mazes we tested were generated with random wall generation, and a sequential process to link nearby walls together. All mazes are presented in Figure S1. Additionally, each maze was slightly stochastic (Figure S2), so that the mazes were larger and more difficult to solve optimally than the chain and the one-step tasks.

## B Robustness to parametrization

Thresholds and parameters can be sensitive to parameter fitting. In this section, we demonstrate that MMRCPD learns accurate models of the environment even when its parameters are wrongly specified. We run experiments on a task of intermediate complexity: the chain task with local volatility. MMRCDP uses the baseline parameters *h* = 10, Δ_*c*_ = 1, Δ_*m*_ = 0.1, max_*mod*_ = 5, and *β* = 3 and is evaluated over 100 seeds. We keep these parameters throughout our analyses and change them one by one to see how they modify the performance of MMRCPD.

**Figure S1:**
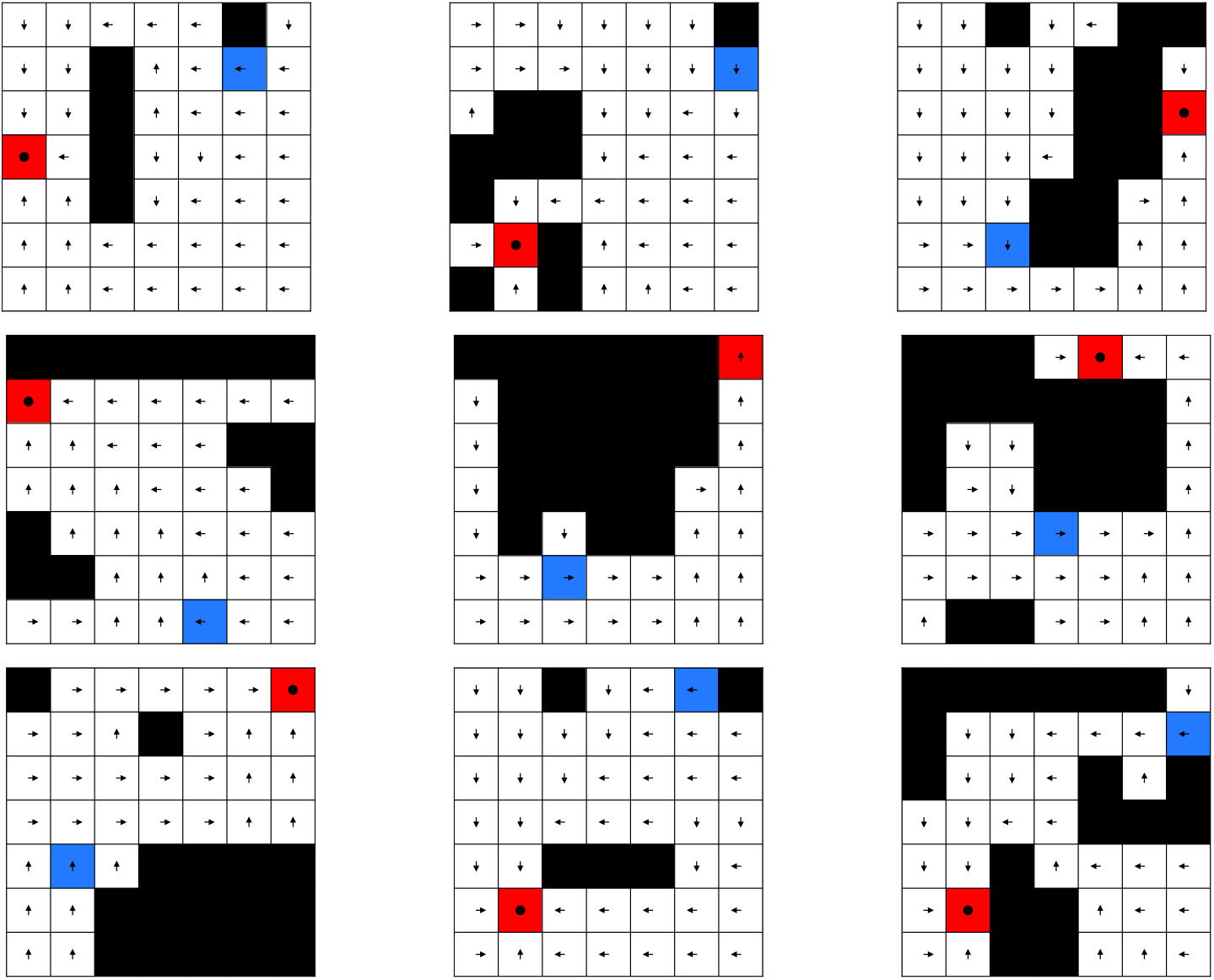
Nine of the ten mazes used in our simulations. The last maze is presented in Figure 6 (Right). These mazes were randomly generated.

**Figure S2:**
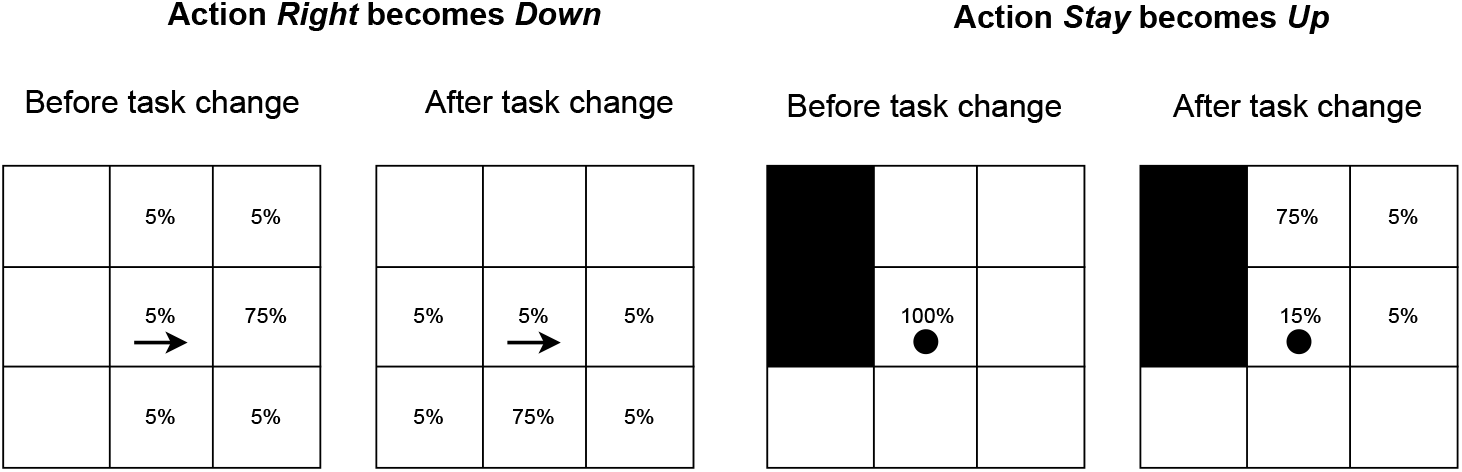
Example of transitions used for the maze environments for a one-step movement action and the *Stay* action. We present the transitions in the absence of walls (Left) and with walls on the left of the position of the agent (Right).

### B.1 Horizon and Model creation

The study horizon *h* of MMRCPD indicates the maximal number of observations the agent uses to evaluate the reliability of its models. If the horizon is too long, the agent may not be able to detect rapid changes. Conversely, a short horizon emphasizes recent observations, potentially causing instability in model creation or model changes. Therefore, the horizon *h* is highly connected to change detection. Another parameter which monitors change detection is the model creation threshold Δ_*c*_. If the model creation threshold is small, the agent might build unnecessary models. If this threshold is high, the agent may not build enough models. If the creation threshold is too high, MMRCPD may behave like an infinite-horizon agent. In deterministic environments, the model creation threshold should be low to create a new model as soon as the agent observes a surprising event. In uncertain environments, the horizon should be high so that the agent samples enough information before deciding whether it should change models.

**Figure S3:**
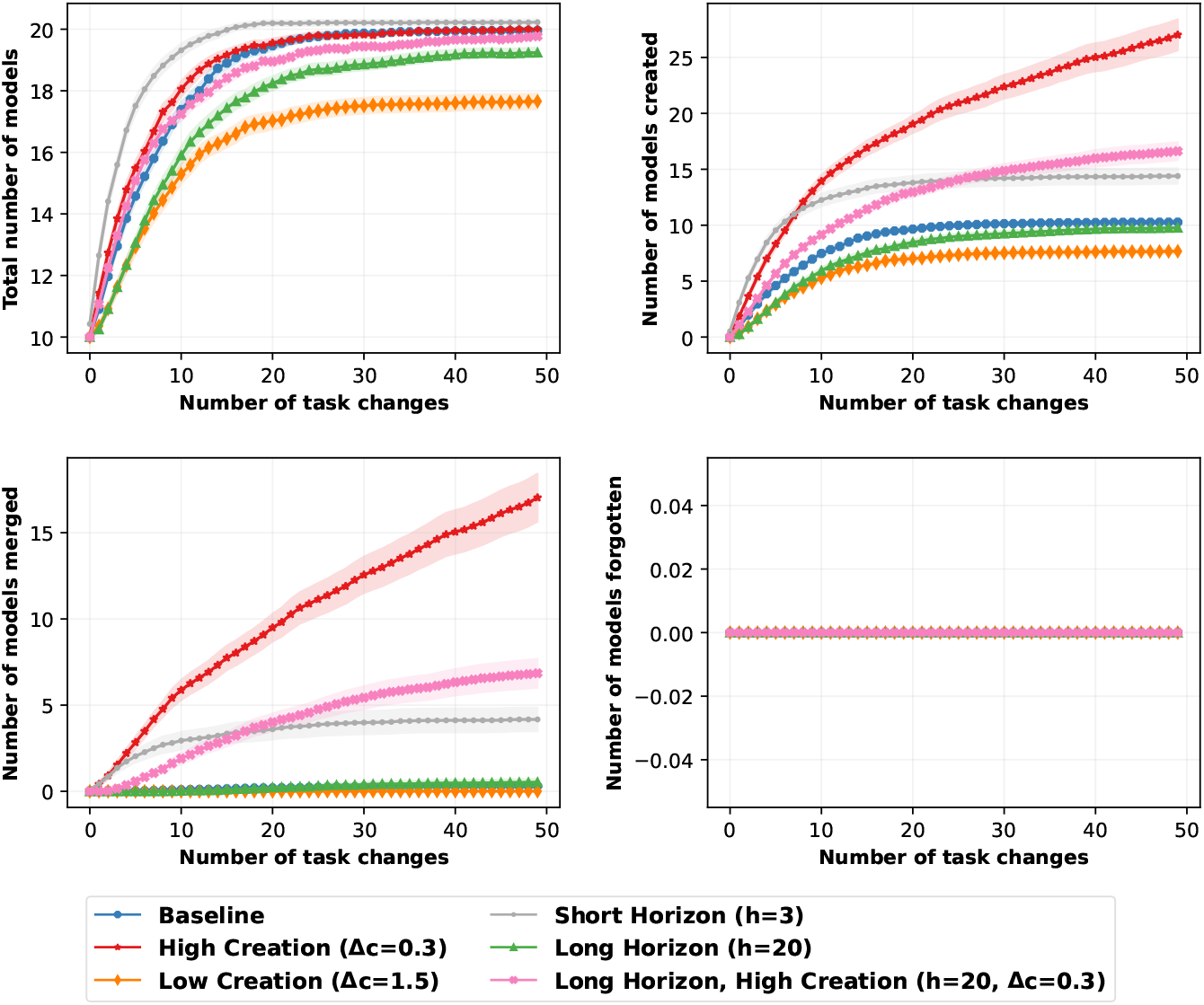
Number of models depending on the horizon and the model creation threshold. (Top-Left) Total number of models. (Top-Right) Number of models created. (Bottom-Left) Number of models merged. (Bottom-Right) Number of models forgotten.

**Figure S4:**
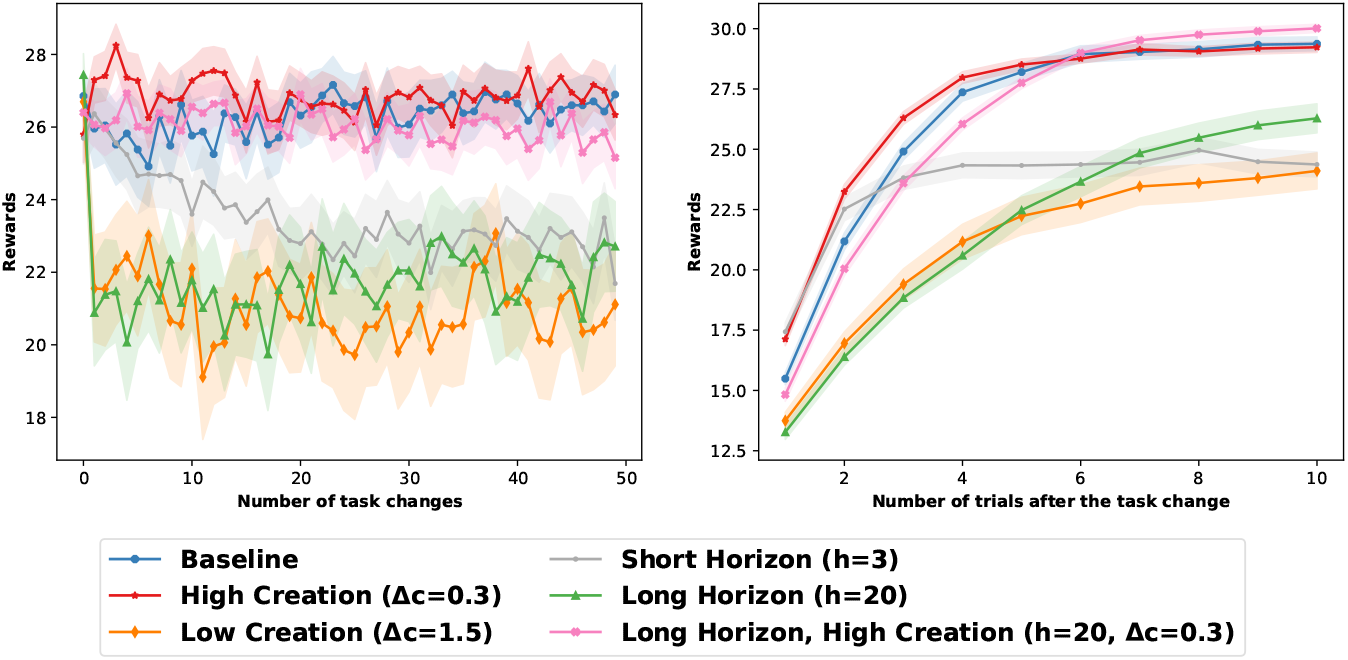
Performance of MMRCPD depending on the horizon and the threshold for model creation.

We compare the evolution of the number of models depending on the horizon *h* and the model creation threshold Δ_*c*_. We study the behavior of an agent with high model creation (Δ_*c*_ = 0.3), an agent with low model creation (Δ_*c*_ = 1.5), an agent with a short horizon (*h* = 3), an agent with a long horizon (*h* = 20) and an agent with a long horizon but a high model creation (*h* = 20, Δ_*c*_ = 0.3).

For all the agents, we plot the total number of models, the number of models created, the number of models merged and the number of models forgotten over time (Figure S3). The baseline agent creates 10 models and stabilizes to 20 total models, which corresponds to the true number of models (2 models for each of the 2 actions for 5 states). It does not merge nor forget models. The agent high creation rate agent (Δ_*c*_ = 0.3) creates more than 25 models throughout the experiment. Nevertheless, the merging module compensates this high creation rate by merging models which are similar. This limits the total number of models to 20 on the studied time scale. The low creation agent (Δ_*c*_ = 1.5) does not create enough models and almost reaches 18 total models on average. The long-horizon agent (*h* = 20) creates models slower than the baseline agent, but almost reaches the appropriate number of models after 50 task changes. The small horizon agent (*h* = 3) creates too many models but stabilizes slightly above 20 models thanks to the merging mechanism. Finally, the agent with a long horizon and high model creation (*h* = 20, Δ_*c*_ = 0.3) almost reaches 20 models.

Although their parameters are incorrectly specified, all the agents created approximately 20 models at the end of the task (between 17 and 21). Yet, the horizon *h* varies between *h* = 3 and *h* = 20 and the model creation threshold Δ_*c*_ varies between Δ_*c*_ = 0.3 and Δ_*c*_ = 1.5. Moreover, agents with high model creation (Δ_*c*_ = 0.3) create and maintain the appropriate number of models thanks to the merging module. These results indicate that the number of models MMRCPD maintains is relatively stable to parameter changes.

Agents with imprecise parameters generally complete the chain task with local volatility (Figure S4). The high-creation agents (Δ_*c*_ = 0.3) gather approximately the same amount of rewards as the baseline agent. The low creation agent (Δ_*c*_ = 1.5) does not create enough models. Therefore, the models it maintains are not accurate and the agent does not attain optimal performance. The short horizon agent (*h* = 3) is fast at detecting task changes but does not perform well when the environment stabilizes. The short horizon agent may be too sensitive to surprising events, changing models too frequently and not maintaining accurate models. The long horizon agent (*h* = 20) takes longer than baseline to detect changes, which reduces its performance on the task.

These results show that, thanks to the merging mechanism, agents which create too many models perform better than agents creating too little models. A high model creation agent can merge similar models, whereas an agent which misses a task change permanently loses information. Creating a model and merging it quickly with a stored model sometimes out-speeds the switching system (for example, with Δ_*c*_ = 0.3 in Figure S4), which only gives the upper hand to a stored model if it explains the latest observations better than the current model.

### B.2 Model Merging and Forgetting

To study how MMRCPD merges and forgets models, we lower the threshold of model creation from Δ_*c*_ = 1 to Δ_*c*_ = 0.3 so that the agent creates too many models. We study the behavior of an agent with high model merging (Δ_*m*_ = 0.3), an agent with no merging (Δ_*m*_ = 0), and an agent with no merging and a low maximal number of models (Δ_*m*_ = 0, max_*mod*_ = 2).

The agent with no merging (Δ_*m*_ = 0) creates too many models and even start forgetting some of them at the end of the experiment (Figure S5). agent with a high merging threshold (Δ_*m*_ = 0.3) merges too many models and does not maintain enough model diversity. This parameter is sensitive as the JS divergence guarantees that the distance between the two distributions is between 0 and log_exp_(2) ≈ 0.7. We voluntarily chose a very low (Δ_*m*_ = 0) and a very high merging threshold (Δ_*m*_ = 0.3) to display these two behaviors. Interpreting the role of the merging parameter is straightforward: the lower it is, the less the agent merges models. The agent with no merging and a low number of maximal models quickly reaches its maximum amount of models (20).

Limiting the number of models may sometimes improve the performance of the agents: the agent with 2 models maximum and no merging performs better than the one with 5 models maximum and no merging (Figure S6). The agent with a high merging threshold performs well, although it does not learn the correct amount of models. This illustrates that agents do not necessarily need very precise models of the environment to perform well when the task is not uncertain.

### B.3 Conclusions on the role of each parameter

The horizon *h* must be long enough to assess model reliability on enough observations. If the horizon is too short and the environment uncertain, the agent may create unnecessary models due to environmental variability. If the horizon is too long, the agent may be slower at detecting changes or miss them out. The larger the model creation threshold Δ_*c*_, the less the agent creates models and the more it performs like a single model infinite-horizon model. If the agent creates too many models because of a wrongly specified horizon or model creation threshold, the merging mechanism limits the total number of models. The higher the merging threshold Δ_*m*_, the more the agent merges models. The maximal number of models max_*mod*_ acts as a safety net, so that the number of models does not increase dramatically when all the other parameters are incorrect. Overall, we illustrated that the parameters we introduced are straightforward and we demonstrated that MMRCPD can perform well even when these parameters are inaccurate.

**Figure S5:**
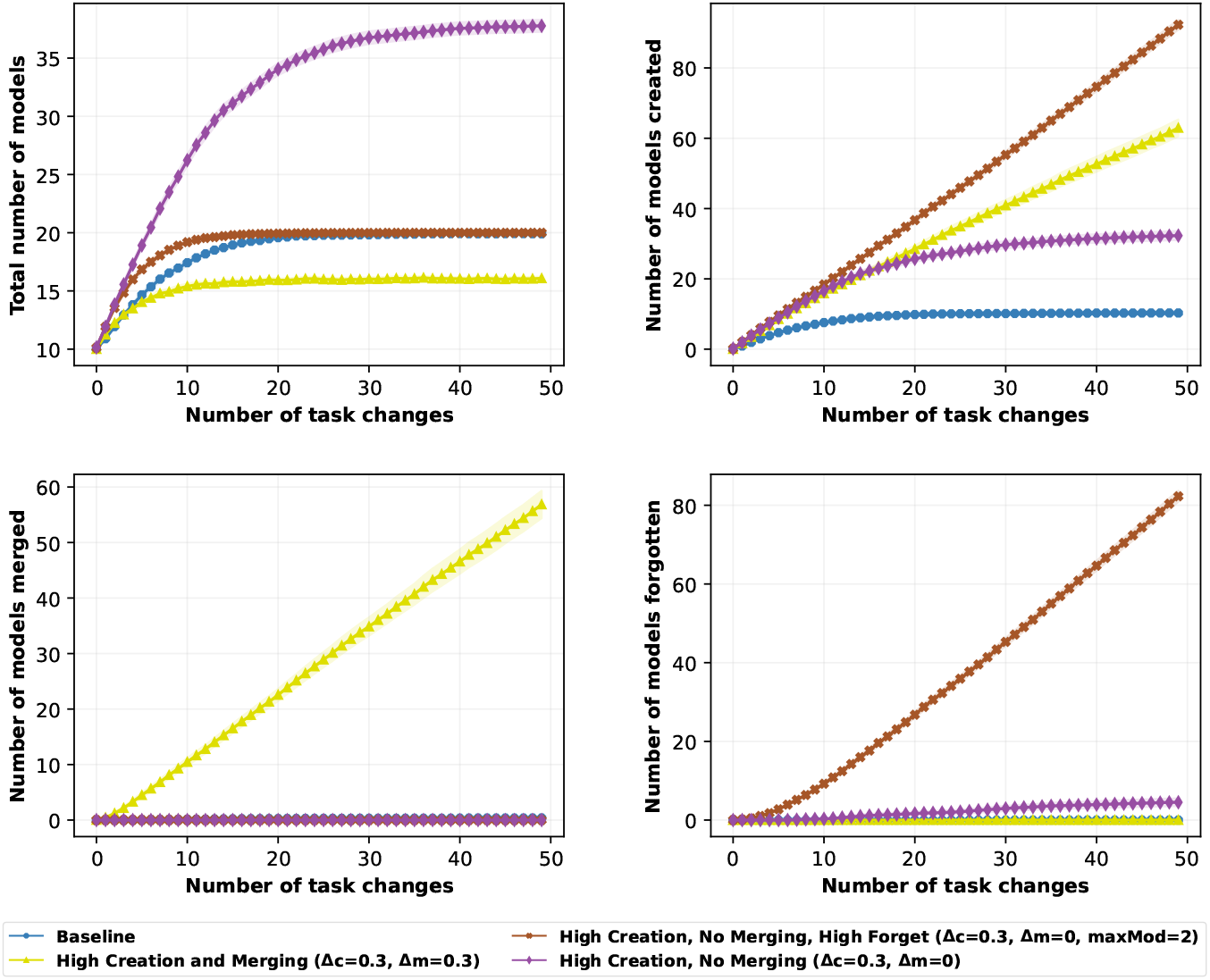
Number of models depending on the model merging threshold and the maximal number of models. (Top-Left) Total number of models. (Top-Right) Number of models created. (Bottom-Left) Number of models merged. (Bottom-Right) Number of models forgotten.

**Figure S6:**
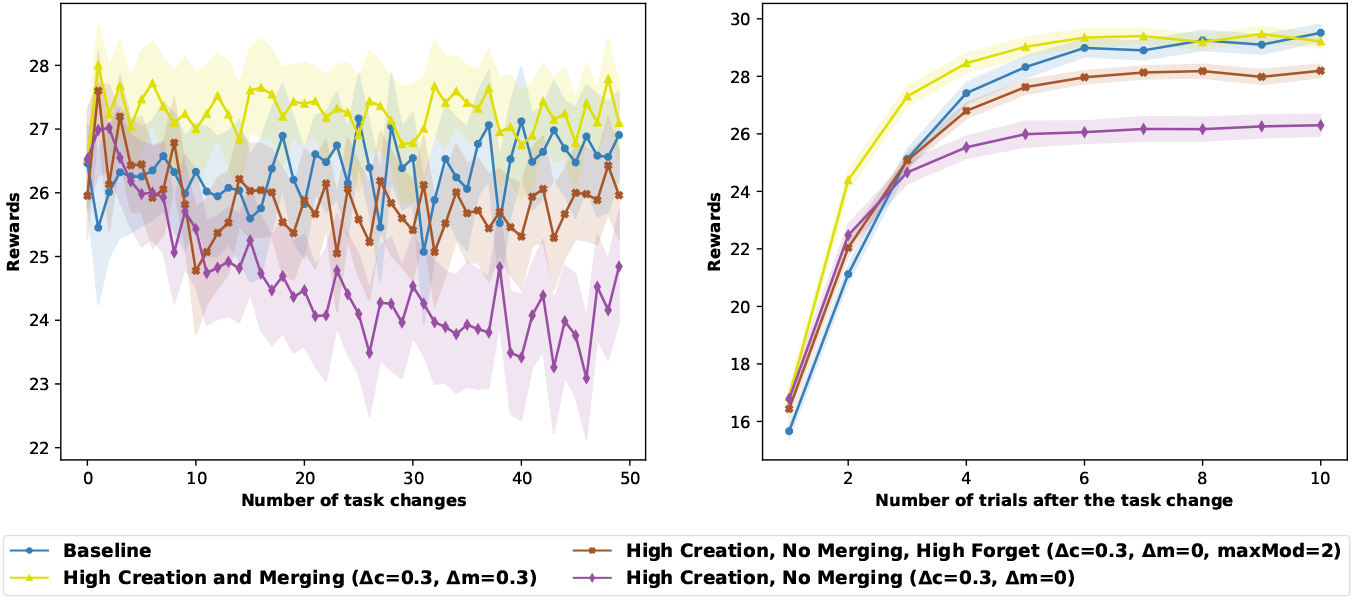
Performance, times and distances to the true model for high creation multi-model agents depending on the merging threshold and the maximal number of models. (Top) Rewards accumulated. (Center) Computational times. (Bottom) Euclidean distance to the true transition model.

## C Computational cost

Agents generally have comparable computational cost on the tasks we simulated. We show the time taken for each agent for the three main classes of environments presented in the main article (Figures S7, S8, and S9).

**Figure S7:**
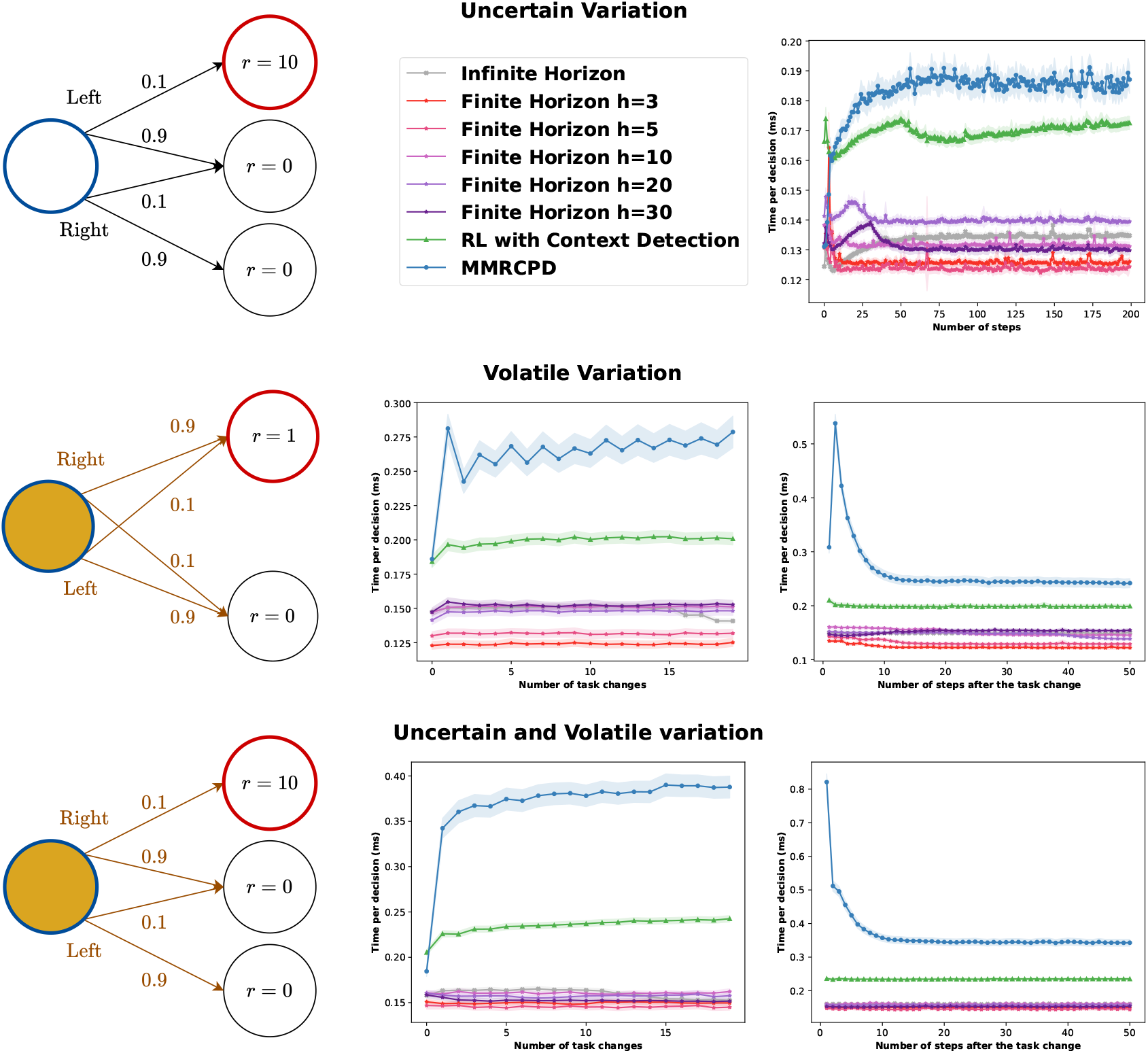
Comparison of performance of the agents in the uncertain (Top), the volatile (Center), and the uncertain and volatile (Bottom) variations of the one-step environment. (Left) Sketch of the environment used. (Right, Center-Center and Center-Bottom) Performance of the agent in the task. For volatile variations, we show two plots: The probability of choosing the best action, averaged over each task change (Left) or averaged over each of the 50 steps between two task changes.

**Figure S8:**
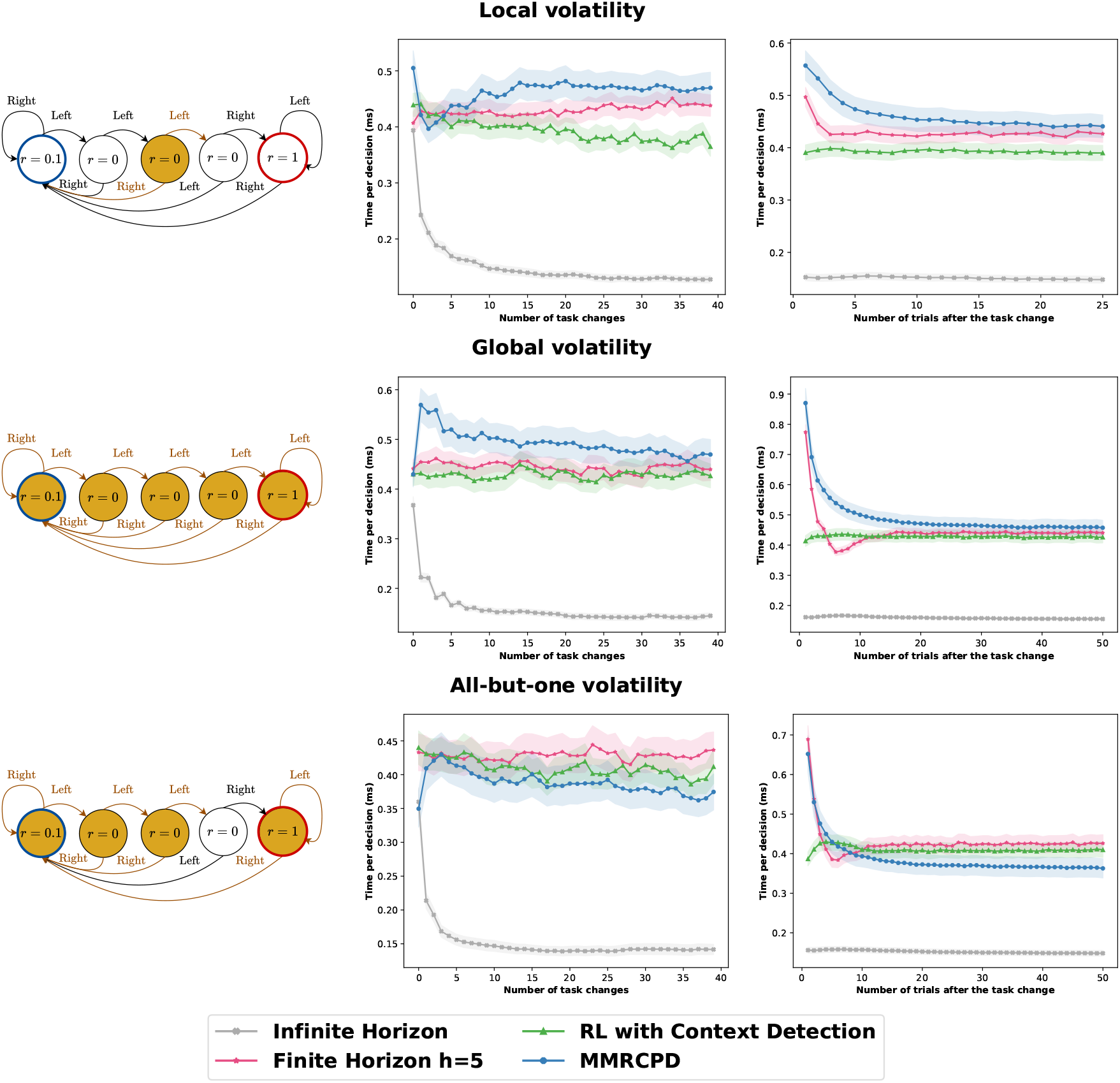
Comparison of performance of local- and context-level approaches based on the task to solve. (Top) When the change is local, context-level method fails to adapt to volatile environments. (Center) When the change is contextual, the context-level method outperforms the local change detection methods, as it detects the task change faster. (Bottom) If the change is global but for one randomly selected state, which changes at each change, context-level methods fail to adapt to changes.

**Figure S9:**
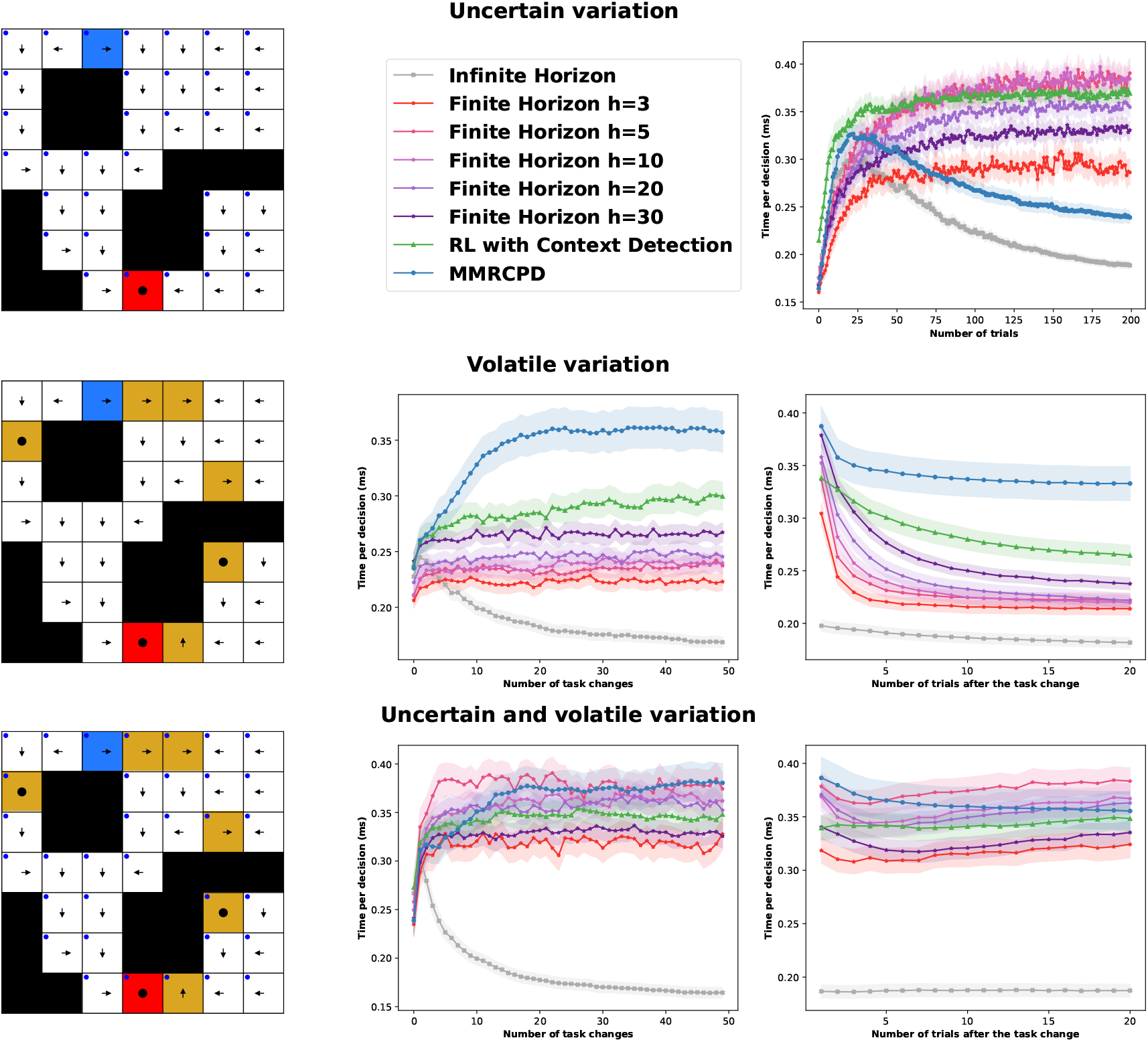
Comparison of performance of the agents in the uncertain (Top), the volatile (Center), and the uncertain and volatile (Bottom) variations of the one-step environment. (Left) Sketch of the environment used. (Right, Center-Center and Center-Bottom) Performance of the agent in the task. For volatile variations, we show two plots: The probability of choosing the best action, averaged over each task change (Left) or averaged over each of the 50 steps between two task changes.

